# The genetic architecture of host response reveals the importance of arbuscular mycorrhizae to maize cultivation

**DOI:** 10.1101/2020.07.06.190223

**Authors:** M. Rosario Ramírez-Flores, Sergio Perez-Limón, Meng Li, Benjamin Barrales-Gamez, Víctor Olalde-Portugal, Ruairidh J. H. Sawers

## Abstract

Arbuscular mycorrhizal fungi (AMF) are ubiquitous in cultivated soils, forming symbiotic relationships with the roots of major crop species. Although studies in controlled conditions have demonstrated the potential of the symbiosis to enhance host plant nutrition and alleviate environmental stress, practical difficulties make it hard to estimate the actual benefit in cultivated fields, not least because of the lack of availability of suitable AMF-free controls. Furthermore, the response can vary depending on the plant variety in a manner which is not fully understood. Here, we implemented a novel strategy based on the selective incorporation of AMF-resistance into a genetic mapping population to evaluate maize response in the field. We found AMF to account for about one third of the grain production in a rain-fed medium input field, as well as to impact the relative performance of plant varieties. Characterization of the genetic architecture of host response allowed us to distinguish mycorrhizal benefit from dependence and indicated a trade-off between mycorrhizal and non-mycorrhizal performance, both at the level of individual QTL and genomewide. This approach is applicable to other crop species, permits further mechanistic analysis and is scalable to full yield trials.

## INTRODUCTION

There is a pressing need to develop sustainable agricultural systems that improve productivity whilst minimizing adverse environmental impacts (Lynch, 2019). In the face of this challenge, there is great interest in the contribution of the arbuscular mycorrhizal (AM) symbiosis and other microbial mutualisms (Sawers et al., 2008; Fester and Sawers, 2011; Bender et al., 2016). AM fungi (AMF) occur broadly in cultivated soils, forming symbiotic relationships with the roots of major crop species and, indeed, many commercial “biofertilizers” are already on the market (Faye et al., 2013). Nonetheless, the value of AM symbiosis in agriculture remains a matter of debate (Ryan and Graham, 2018; Rillig et al., 2019). Experimental studies under controlled conditions have demonstrated that AMF have much to offer host plants, including enhanced uptake of immobile soil nutrients (especially phosphorus) and water, and improved tolerance to a range of abiotic and biotic stresses (Borowicz 2001; Chandrasekaran et al., 2014; Augé et al., 2015; Chiu and Paszkowski, 2019). AMF also contribute to important ecosystem services (van der Heijden et al., 2015), including promoting soil aggregation (Rillig and Mummey, 2006) and reducing nutrient loss through leaching and emission (Bender et al., 2014; Cavagnaro et al., 2015). However, efforts to integrate AMF into agricultural practice are hampered by the difficulty of evaluating host response in the field and the lack of appropriate AMF free controls: when exogenous fungal inoculum is applied to a field, it is difficult to gauge its effectiveness with respect to the native fungal community; when looking to evaluate the impact of the native AMF community under any given agronomic scenario, there is no easy way to estimate what the baseline performance of plants would be if the AMF were not present. Although experimental designs using chemical treatments and/or mechanical barriers might be implemented at small scale they are limited and not practical for use in larger genetic studies or trials. Here, to address these difficulties, we propose the use of genetic variants conferring resistance to AMF in the design of experiments to study AM symbiosis in the field.

AM symbiosis dates back over 450 million years (Strullu-Derrien et al., 2018), In contrast, agriculture is a recent development. Modern breeding efforts have focused on maximizing yield under homogeneous environments with large quantities of synthetic inputs, and it is not clear how the breeding process has impacted the AM symbiosis (discussed in (Sawers et al., 2008; Sawers et al., 2018). Nonetheless, modern varieties of our major crops do retain the capacity to interact with AMF and the molecular machinery necessary for symbiotic establishment (*e.g.* Gutjahr et al., 2008). Elegant molecular genetic analyses, largely conducted in *Medicago truncatula* and *Lotus japonicus*, have identified a series of plant-encoded receptor and signal-transduction components necessary for AM symbiosis and, in relevant host plants, for interaction with nodule-forming rhizobia. Fungal signals are perceived by the host SYMRK (DMI2) receptor, triggering cellular calcium spiking events through the action of the nuclear ion-channels CASTOR and POLLUX (DMI1) and activate downstream events via CCamK (DMI3) and CYCLOPS (IPD3) transduction components (Parniske, 2008; Zipfel and Oldroyd, 2017). Notwithstanding the great antiquity of AM symbiosis, there is little redundancy in symbiotic signalling and single gene mutations in the host can be sufficient to generate a complete early block in symbiotic interaction (*e.g.* Charpentier et al., 2008; Gutjahr et al., 2008). In non-nodulating hosts, mutations in symbiotic signaling pathway genes appear to specifically impact AM symbiosis and do not condition obvious pleiotropic effects on plant development. To date, of the major cereal crops, mutations in the symbiotic signalling pathway have only been reported in rice ((Charpentier et al., 2008; Gutjahr et al., 2008)), although the genes themselves can be unambiguously identified in the genomes of plants such as maize, wheat and sorghum.

While the symbiotic pathway is essential for plant interaction with AMF, there is no evidence that it plays a role in regulating the progression of the symbiosis or that variation in the pathway underlies the range of host response that is observed among different plant varieties (Kaeppler et al., 2000; Mascher et al., 2014; Sawers et al., 2017; Watts-Williams et al., 2019). Indeed, defining variation in plant response to AMF can in itself be challenging (discussed in Janos, 2007; Sawers et al., 2010). Here, we define *response* as the difference between mycorrhizal and non-mycorrhizal plants, applied to any phenotypic measurement. In addition, we define *dependence* as the capacity of a given variety to perform in the absence of AMF (Janos, 2007), and *benefit* as the degree to which a plant host profits from the association. Crucially, variation in host response confounds differences in both dependence and benefit. The optimization of AM symbiosis in crop varieties is concerned with benefit: there is no interest in elevating host response by increasing dependence (Sawers et al., 2008). However, these somewhat abstract concepts can be difficult to distinguish, especially when comparing a small number of varieties at the whole genotype level: if one variety shows a greater proportional increase in growth when inoculated with AMF than another, does this reflect a superior ability to profit from the symbiotic interaction, or poorer performance in the absence of symbiosis? This question has relevance both to hypotheses concerning the impact of past selection of AM symbiosis and to any future breeding effort to optimize the interaction. Mechanistic understanding can provide useful insight: *e.g.* response variation linked to differences in root development might be reasonably assigned to dependence (Paszkowski and Boller, 2002), response variation linked to differences in the growth of root-external fungal hyphae to benefit (Sawers et al., 2017). However, it is not practical to characterize all the physiological differences among a larger collection of varieties and far from trivial to use these data to interpret variation in host response to AMF. Here, we propose to distinguish dependence and benefit using the concepts of genotype x environment interaction (Des Marais and Juenger, 2010), where the mycorrhizal and non-mycorrhizal conditions are considered as two distinct “environments”. In conjunction with the ability to perform large scale mapping experiments, the GEI framework has the potential to define dependence and response at the level of the variety and at the level of the action of individual genomic regions.

In this study, we identify a loss-of-function allele of the maize *CASTOR* ortholog, confirm it confers resistance to AMF and selectively incorporate it into a biparental population for quantitative trait locus (QTL) mapping. The resulting population contained both AM susceptible and resistant families, allowing us to map QTL for agronomically important yield components in mycorrhizal and non-mycorrhizal “environments”. Our results indicate AM symbiosis tp makes a substantial contribution to maize performance in the field. Furthermore, we present an empirical demonstration of the dependence and benefit variation, as well as evidence of a trade-off between the two.

## RESULTS

### Mutation in a maize common symbiosis gene demonstrates the importance of arbuscular mycorrhizae in the field

To generate AMF-resistant maize varieties for use as a control in field experiments, we identified maize orthologs of the common-symbiosis genes *CASTOR* and *POLLUX* (*DMI1*) (Parniske, 2008). *CASTOR* and *POLLUX* were found to be single copy genes in maize, and designated *Hunahpu* (*Hun1*; GRMZM2G099160; Zm00001d012863) and *Xbalanque* (*Xba1*; GRMZM2G110897; Zm00001d042694) after the hero twins of Mayan mythology (Ramírez-Flores, 2015). We searched publicly available genetic resources (McCarty et al., 2005) and identified a transposon-insertion allele of the *Hun1* gene (*hun1-2::mu*, hereafter, *hun1*) that we found to confer resistance to mycorrhizal colonization (Ramírez-Flores, 2015). We then crossed the *hun1* mutant in its original temperate W22 background to the subtropical inbred line CML312 and advanced material to generate a mapping population of 73 wild-type (susceptible; *AMF-S*) and 64 *hun1* (resistant; *AMF-R*) F_2:3_ families (Fig. 1A, S2). We confirmed that AM resistance was stable in the field, colonization being absent or greatly reduced in *AMF-R* families in a preliminary trial (Fig. S3). In a second preliminary experiment, we grew *AMF-S* and *AMF-R* families under highly managed nutrient-replete field conditions and found there to be no major difference in performance, indicating that *hun1* does not condition a general defect in plant growth or show marked pleiotropic effects beyond the block in mycorrhizal colonization (Fig. S4, S5).

**Figure 1.**
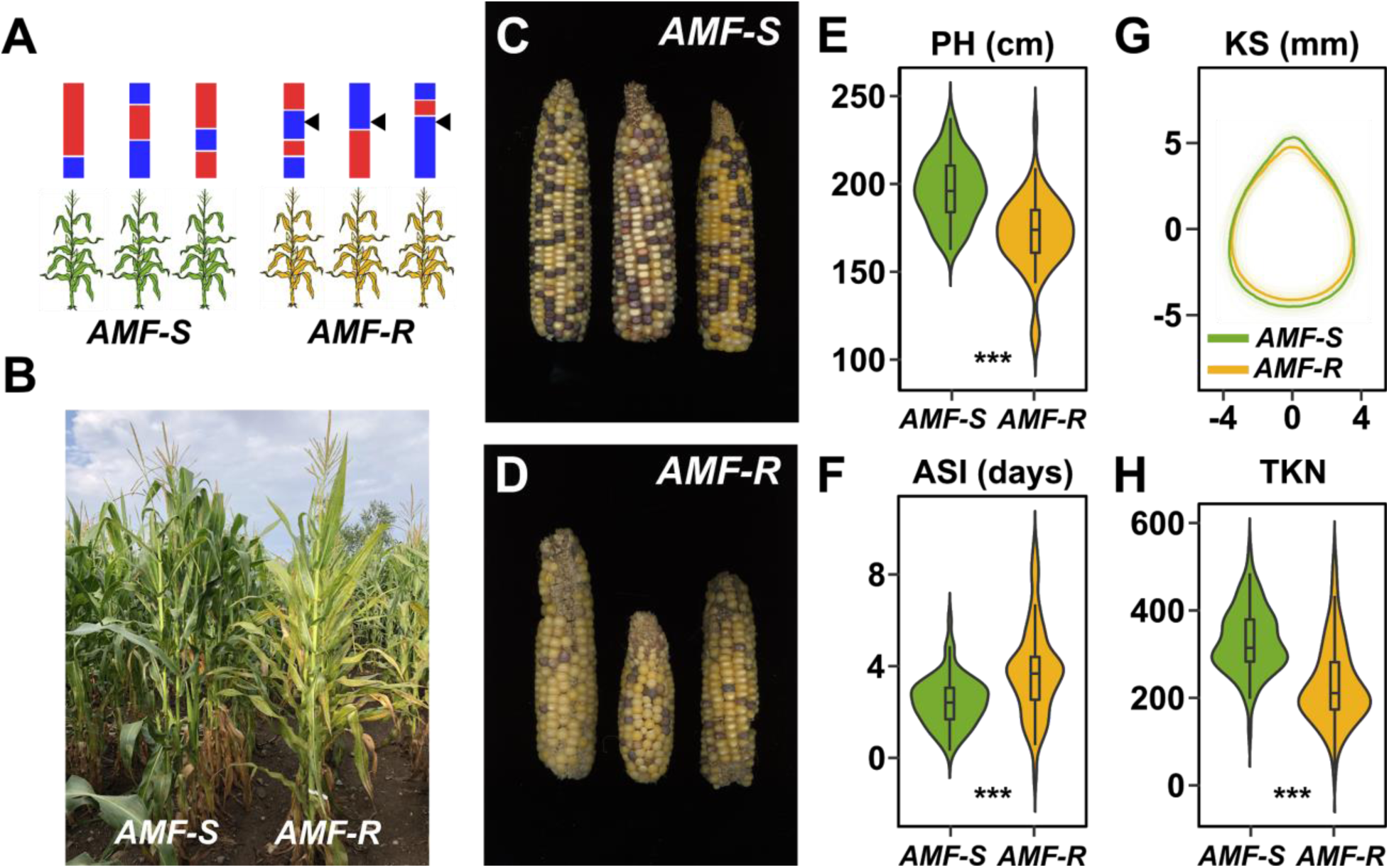
Mycorrhizal symbiosis impacts maize growth under cultivation. **A**, Susceptible (*AMF-S*) and resistant (*AMF-R*) families segregate genomic content from two founder parents, shown as red and blue bars. *AMF-R* families are homozygous for the *hun1-2* mutation, blocking AM symbiosis (black arrows). **B**, Border between representative *AMF-S* and *AMF-R* plots, Jalisco, Mexico, 2019. **C**, Representative AMF-S ears. **D**, Representative AMF-R ears. **E**, Plant height (PH) of 73 *AMF-S* and 64 *AMF-R* families. **F**, Kernel shape (KS). **G**, Anthesis-silking interval (ASI). **H**, Total kernel number (TKN). The violin plots in **E, F** and **H** are a hybrid of boxplot and density plot. The box represents the interquartile range with the horizontal line representing the median and whiskers representing 1.5 times the interquartile ranges. The shape of the violin plot represents probability density of data at different values along the y-axis.

To estimate the importance of AM symbiosis, we evaluated *AMF-S* and *AMF-R* families in a replicated trial under rain-fed, medium-input, conditions in Ameca, Mexico (Fig. 1A, S4, S6; triplicated three-row plots, ~1,200 rows of 15 individuals for a total of ~ 20, 000 plants). We observed *AMF-R* families to generally exhibit mild-chlorosis (Fig. 1B, S7) and to produce visibly poorer ears than *AMF-S* families (Fig. 1C-D). We collected fifteen phenological and morphological traits and yield-components (Fig. 1E-H, S8; Table S1), eleven of which showed a significant marginal difference between *AMF-S* and *AMF-R* families, equating to host response ranging from 3 to 51 % (Table 1). *AMF-R* families were reduced in stature and delayed in silking (female flowering), resulting in an extension of anthesis-silking interval (ASI; Fig 1E, F; Table S1) - a classic symptom of abiotic stress in maize. Ear size was reduced in *AMF-R* families, along with ear and total kernel weight (Table 1; Fig. S9). Individual kernels were not significantly different in weight or size between *AMF-S* and *AMF-R* families (Fig. 1G; Table 1). The total number of kernels per ear, however, was greatly reduced in *AMF-R* families (Fig. 1H; Table 1), indicative of poor seed set, a possible consequence of increased ASI.

**Table 1.**
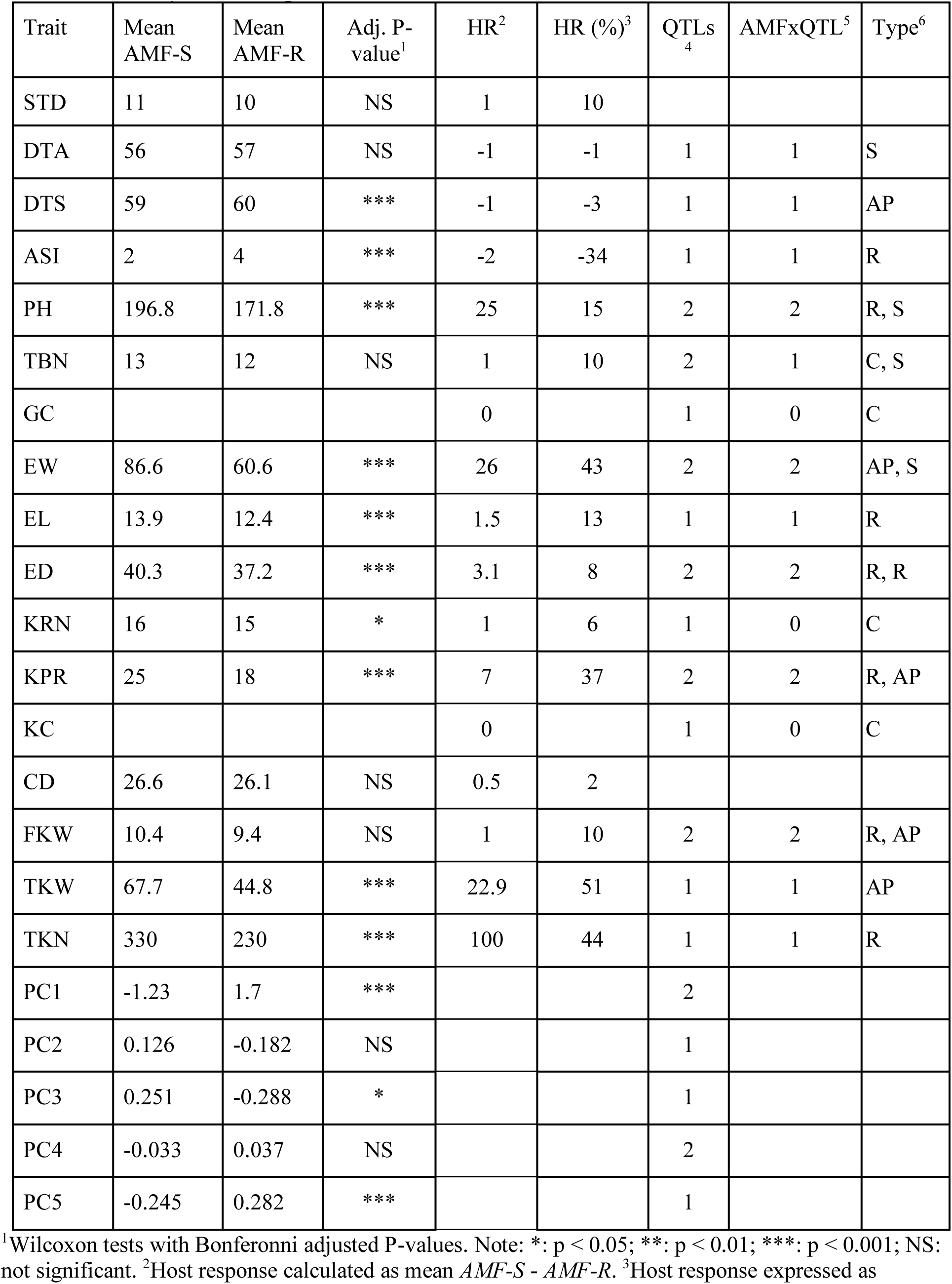
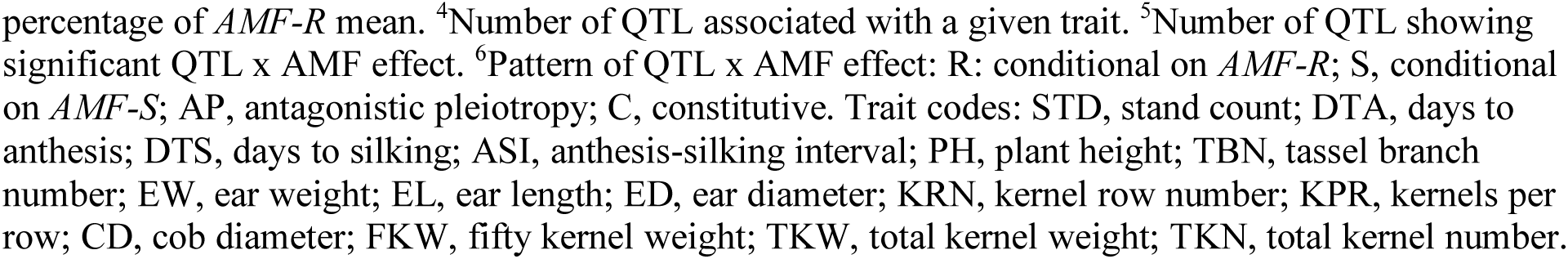
Summary of host response and QTLs

| Trait | Mean AMF-S | Mean AMF-R | Adj. P-value<sup>1</sup> | HR<sup>2</sup> | HR (%)<sup>3</sup> | QTLs<sup>4</sup> | AMFxQTL<sup>5</sup> | Type<sup>6</sup> |
| --- | --- | --- | --- | --- | --- | --- | --- | --- |
| STD | 11 | 10 | NS | 1 | 10 |  |  |  |
| DTA | 56 | 57 | NS | -1 | -1 | 1 | 1 | S |
| DTS | 59 | 60 | *** | -1 | -3 | 1 | 1 | AP |
| ASI | 2 | 4 | *** | -2 | -34 | 1 | 1 | R |
| PH | 196.8 | 171.8 | *** | 25 | 15 | 2 | 2 | R, S |
| TBN | 13 | 12 | NS | 1 | 10 | 2 | 1 | C, S |
| GC |  |  |  | 0 |  | 1 | 0 | C |
| EW | 86.6 | 60.6 | *** | 26 | 43 | 2 | 2 | AP, S |
| EL | 13.9 | 12.4 | *** | 1.5 | 13 | 1 | 1 | R |
| ED | 40.3 | 37.2 | *** | 3.1 | 8 | 2 | 2 | R, R |
| KRN | 16 | 15 | * | 1 | 6 | 1 | 0 | C |
| KPR | 25 | 18 | *** | 7 | 37 | 2 | 2 | R, AP |
| KC |  |  |  | 0 |  | 1 | 0 | C |
| CD | 26.6 | 26.1 | NS | 0.5 | 2 |  |  |  |
| FKW | 10.4 | 9.4 | NS | 1 | 10 | 2 | 2 | R, AP |
| TKW | 67.7 | 44.8 | *** | 22.9 | 51 | 1 | 1 | AP |
| TKN | 330 | 230 | *** | 100 | 44 | 1 | 1 | R |
| PC1 | -1.23 | 1.7 | *** |  |  | 2 |  |  |
| PC2 | 0.126 | -0.182 | NS |  |  | 1 |  |  |
| PC3 | 0.251 | -0.288 | * |  |  | 1 |  |  |
| PC4 | -0.033 | 0.037 | NS |  |  | 2 |  |  |
| PC5 | -0.245 | 0.282 | *** |  |  | 1 |  |  |

<sup>1</sup>Wilcoxon tests with Bonferonni adjusted P-values. Note: \*: p < 0.05; \*\*: p < 0.01; \*\*\*: p < 0.001; NS: not significant. <sup>2</sup>Host response calculated as mean *AMF-S* - *AMF-R*. <sup>3</sup>Host response expressed as
percentage of *AMF-R* mean. <sup>4</sup>Number of QTL associated with a given trait. <sup>5</sup>Number of QTL showing significant QTL x AMF effect. <sup>6</sup>Pattern of QTL x AMF effect: R: conditional on *AMF-R*; S, conditional on *AMF-S*; AP, antagonistic pleiotropy; C, constitutive. Trait codes: STD, stand count; DTA, days to anthesis; DTS, days to silking; ASI, anthesis-silking interval; PH, plant height; TBN, tassel branch number; EW, ear weight; EL, ear length; ED, ear diameter; KRN, kernel row number; KPR, kernels per row; CD, cob diameter; FKW, fifty kernel weight; TKW, total kernel weight; TKN, total kernel number.

### QTL x AMF effects underlie variation in mycorrhiza response

To characterize the genetic architecture of host response, we performed a Quantitative Trait Locus (QTL) analysis using the trait estimates obtained for the F_2:3_ families and the genotypes of their respective F_2_ parents (Fig. S10, S11). We combined *AMF-S* and *AMF-R* families in a single analysis, treating the *Hun* genotype as an interactive covariate. Under this model, the *Hun* additive effect (hereafter, AMF effect) estimated the marginal host response across all families, QTL additive effects captured genetic differences between the parents, and QTL x AMF effects indicated underlying heritable differences in the response of CML312 and W22 parents. We identified a total of 24 QTLs, 17 of which also showed AMF x QTL interaction (Fig. 2A, S12; Table 1). We combined significant QTL, AMF and AMF x QTL terms, into a single multiple-QTL model for each trait and calculated the percentage of phenotypic variation explained by AMF, by additive QTL and by AMF x QTL terms (Fig. 2B; Table S2). For plant height, ear weight and total kernel weight, the combination of AMF and AMF x QTL effects explained more than half of the total genetic variation (based on estimation of H^2^), and over a quarter of the total phenotypic variance (Fig. 2B). The identification of significant AMF x QTL effects reveals genetic variation for host response between the CML312 and W22 parents.

**Figure 2.**
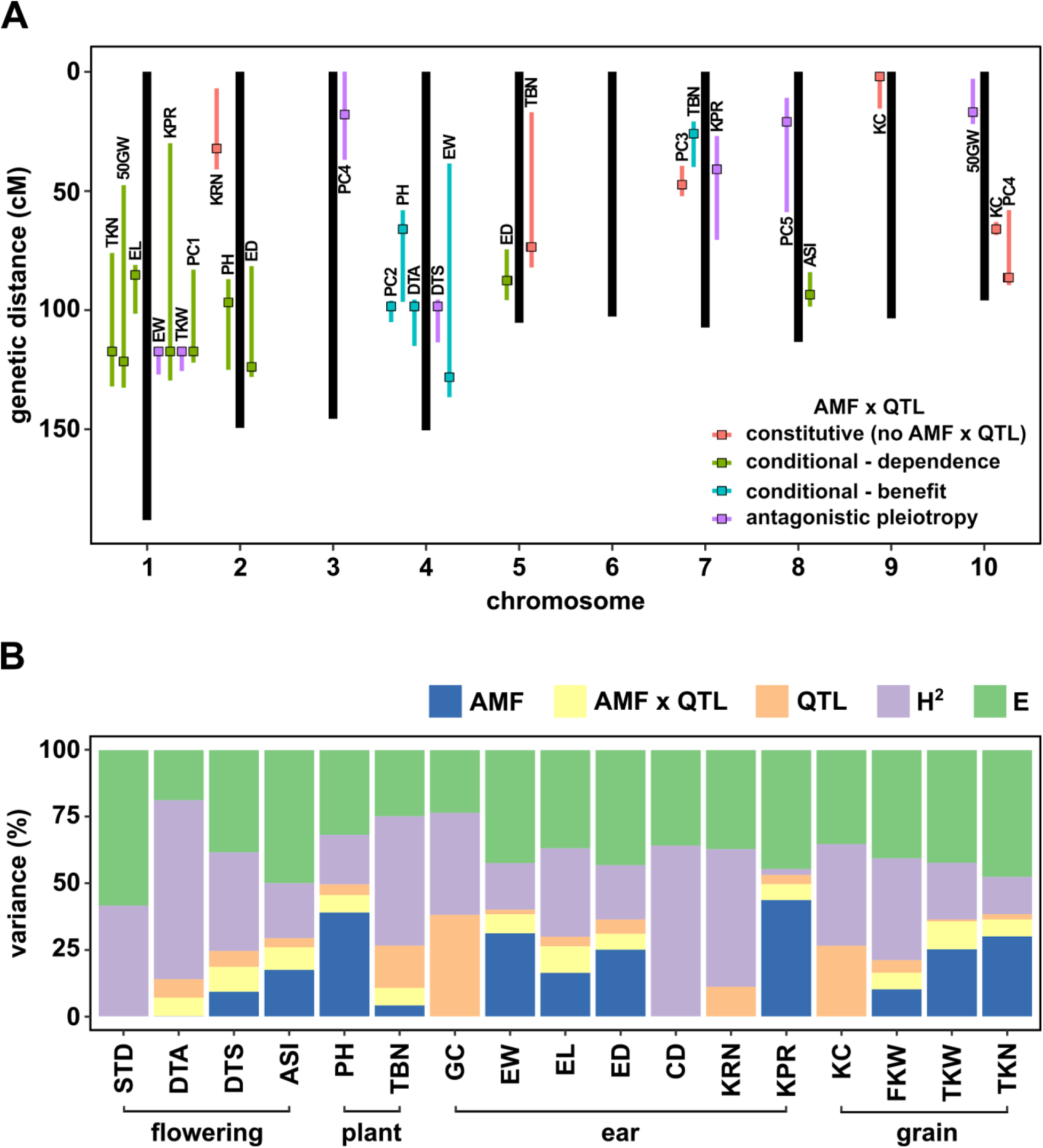
QTL x AMF effects contribute significantly to variation in host response. **A)** Genomic position of QTL identified in this study. Boxes indicate the position of the peak marker. Bars represent the drop-1 LOD interval. Colors denote patterns of AMF x QTL interaction as described in the text. **B)** The contribution of different components to phenotypic variance in different plant traits among *AMF-S* and *AMF-R* families. Total genetic variance (H^2^) was estimated based on differences between experimental blocks. Variance explained by the QTL model for each trait is partitioned into that explained by the additive effect of AMF, the additional variation explained by interactive QTL and QTL x AMF interaction (QTL x AMF) and the additional variation explained by non-additive QTL. Traits codes as in Table 1.

### The genetic architecture of response variation distinguishes dependence and benefit

Variation in host response variation confounds differences in dependence and benefit (Fig. 3A, B, S13; Janos, 2007; Sawers et al., 2010) - the former being the capacity of a given variety to perform in the absence of AMF, the latter the degree to which a plant host profits from the association. Having identified significant AMF x QTL interaction, we distinguished dependence and benefit by estimating the size of the QTL effect separately for *AMF-S* and *AMF-R* families (Fig. S14). For 12 of the 17 QTLs showing significant interaction, the effect was *conditional* - *i.e.* expressed in either the *AMF-S* or *AMF-R* families, but not in both (Fig. 2A, 3; Table 1; Des Marais and Juenger, 2010). We considered conditional QTLs expressed specifically in the *AMF-R* families to represent variation in dependence. Conversely, we considered conditional QTLs expressed specifically in the *AMF-S* families to be variation in benefit (Fig. 2, 3; Table 1). Two regions of the genome were associated with multiple QTLs, suggesting a shared mechanistic basis: a region on the long arm of chromosome (chr) 1 linked to ear-traits and a region on chr 4 linked to plant height and flowering time (Fig. 2A). Combining traits using Principal Components (PCs) further refined these two QTL hot-spots and clearly demonstrated the difference between dependence and benefit (Fig. 3D, S15. Table 1).

**Figure 3.**
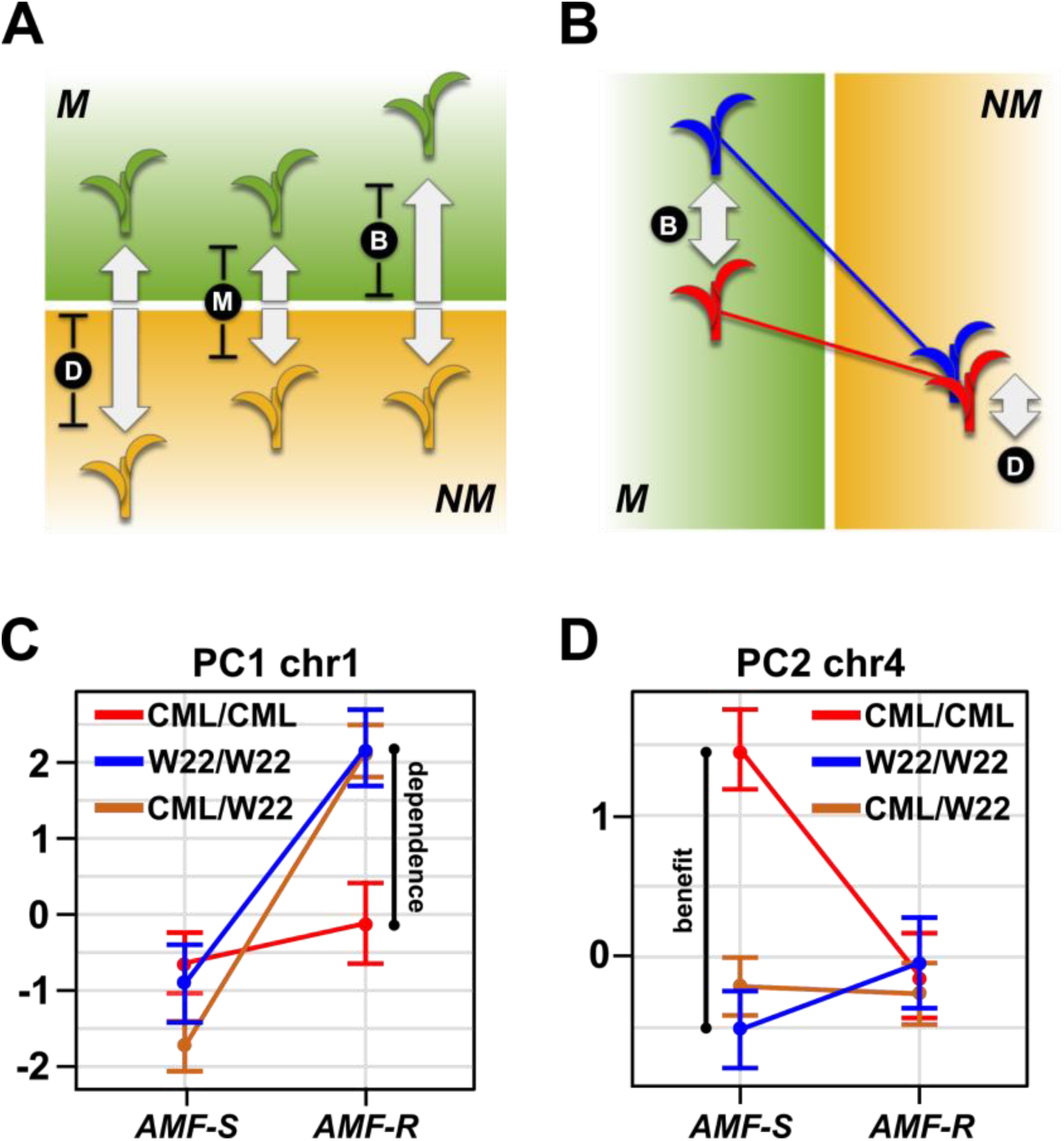
Mycorrhiza response confounds benefit and dependence. **A**, host response (R) is the difference in trait value between mycorrhizal (M; green) and non-mycorrhizal (NM, yellow) plants. Increased response can result from either greater dependence (D; poorer performance of NM plants) or greater benefit (B; greater performance of M plants). **B**, QTL x AMF effects underlying variation in response reveal the balance of benefit and dependence. The difference in trait value between plants carrying the CML312 (red) or W22 (blue) allele. In this theoretical example, the effect is conditional on mycorrhizal colonization, reflecting a difference in benefit more than dependence. **C, D**, effect plots for major QTL associated with PC1 and PC2, respectively. The PC1 QTL is conditional on *AMF-R*, indicating a difference in dependence. The PC2 QTL is conditional on *AMF-S*, indicating a difference in benefit.

### Antagonistic QTL effects suggest a trade-off between mycorrhizal and non-mycorrhizal growth

Several QTLs showed a more extreme form of QTL x AMF in which the effect was expressed in both *AMF-S* and *AMF-R* populations, but with a change of sign (*i.e.* a “swap” in the relative performance of the parental alleles) - a condition described as *antagonistic pleiotropy* (Fig. 4A, S13, S14; Des Marais and Juenger, 2010). Under such genetic architecture, the superior allele for mycorrhizal plants is detrimental to performance in the absence of mycorrhiza, and *vice versa*, providing evidence for a trade-off at the single-locus level. Significantly, several QTLs associated with key yield components showed antagonistic pleiotropy (Fig. 2, 4B, 4C, S14; Table 1). Analogous “rank changing” effects were also seen at the level of the whole genotype. Although the mapping design did not permit direct evaluation of the same genotype as both *AMF-S* and *AMF-R*, we could fit QTL models to all genotypes across both levels of AMF. The resulting estimates showed evidence of rank change with respect to performance with or without AMF (Fig. 4D). To further explore response at the genotype level, we used whole genome models to predict mycorrhizal and non-mycorrhizal trait values across the 137 families. Specifically, we used the *AMF-S* families to train a model for mycorrhizal performance that was then applied across the population. Similarly, we used the *AMF-R* families to train a model for non-mycorrhizal performance. Whole genome predictions aligned well with the observed values of their training sets, although they did not capture the more extreme observed values (Fig. 4E). Comparison of the two models indicated that genotypes associated with the highest values of the major yield components in one condition were unexceptional or poor in the other (Fig. 4F), indicating trade-off between dependence and benefit.

**Figure 4.**
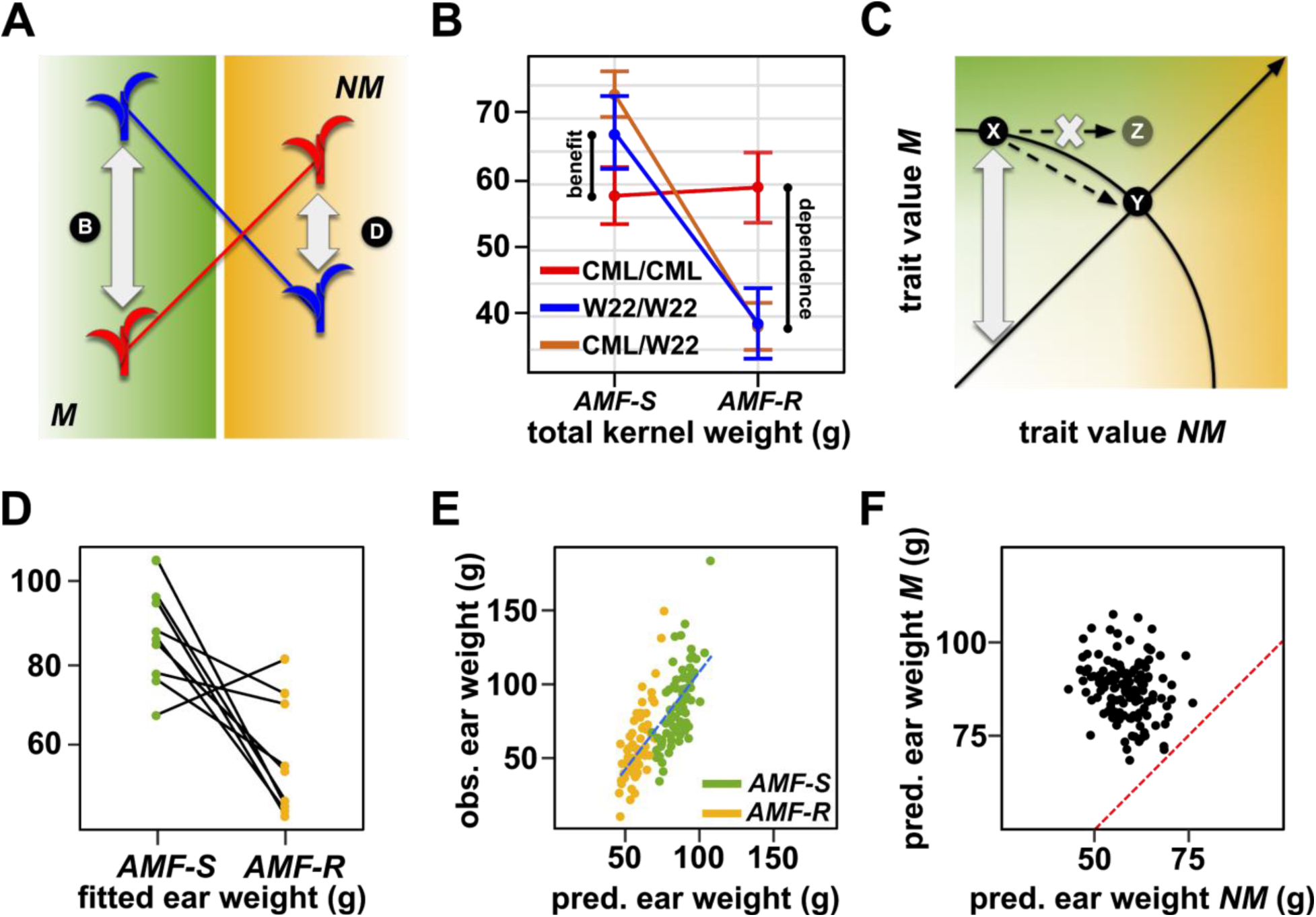
The genetic architecture of mycorrhiza response implies a trade-off between dependence and benefit. **A**, QTL x AMF effects underlying variation in response reveal the balance of benefit and dependence. The difference in trait value between plants carrying the CML312 (red) or W22 (blue) allele. In this theoretical example, the effect is conditional on mycorrhizal colonization, reflecting a difference in benefit more than dependence. **B**, QTL effects for a total kernel weight (TKW) QTL located on Chromosome 1. TKW of families homozygous for the CML312 allele at this locus (red) is stable across susceptible (*AMF-S*) and resistant (*AMF-R*) families. In contrast, families homozygous for the W22 allele (blue) or segregating at this locus (brown), show superior TKW if *AMF-S* but lower TKW if *AMF-R*. In this extreme example of QTL x AMF, the QTL is linked to differences in both dependence and benefit, an example of antagonistic pleiotropy or “trade-off”. **C**, Theoretical trade-off between the performance of non-mycorrhizal (NM) and mycorrhizal (M) plants. For any position on this plot, the vertical distance above the diagonal indicates the magnitude of host response. Trade-off prevents occupancy of the upper-right of the plot, defining a so-called “Pareto front” (solid arc). Here, an increase in the NM performance of variety X is necessarily accompanied by a reduction in M performance, as described by movement along the arc to condition Y. Biological constraint prevents the path towards condition Z which would represent NM improvement without any negative impact on M. **D**, Fitted ear weight values for all genotypes from the QTL model (2 QTL and their interactions with AMF) under both levels of AMF (*AMF-S* and *AMF-R*). Line segments connect the same QTL genotype under the two AMF levels. **E**, Observed ear weight against whole-genome prediction for 73 susceptible (*AMF-S*) and 64 resistant (*AMF-R*) families. Best fit regression line in dashed blue. **F**, Predicted ear weights for all 137 genotypes based on *AMF-S* (M) and *AMF-R* (NC) whole-genome models. Dashed red line shows *AMF-S* = *AMF-R*, *i.e.* no host response.

## DISCUSSION

We have presented evidence that AM symbiosis makes a significant contribution to maize performance. Extrapolating our observations to a cultivated field planted at 80,000 plants/ha (typical in the locality of the trial site), we estimate that AM colonization would contribute ~ 2 tonnes/ha to a total yield of ~ 5.5 tonnes/ha. Although a rough figure to be treated with caution, this estimate serves to provide a striking indication of the importance of AMF, and is consistent with the previous evaluation of an inbred mutant line that linked loss of mycorrhizal phosphate uptake with a 15% reduction in ear-weight (Willmann et al., 2013). Although maize varieties have consistently been shown to benefit from AM symbiosis (*e.g.* Kaeppler et al., 2000; Sawers et al., 2017), the impact of AMF in other plant species may vary along a continuum from mutualism to parasitism (Bergmann et al., 2020; Johnson et al., 1997). In wild species, host response is centered around neutral commensalistic interactions, with equal occurrence of parasitism and mutualism (Klironomos, 2003). A similar range of host response has also been reported in crop plants (Wen et al., 2019). In sorghum, a close relative of maize, host response varies from positive to negative depending on both the plant variety and the AMF species (Ramírez-Flores *et al*., unpublished observations; Watts-Williams et al., 2019). However, host response has typically been measured in young plants, grown under controlled conditions, and the values obtained may under-estimate the true importance of AM symbiosis in the field.

We found evidence for substantial variation in host response among plant genotypes. Based on the range of fitted values obtained from the total grain weight multiple-QTL model, we predict a range of yield response from 0.4 to 3.7 tonnes/ha at 80,000 plants/ha. In terms of variance explained, AMF x QTL effects were more important than the QTL main effects for total kernel weight, total kernel number and fifty-kernel weight. While we saw no evidence that our W22 and CML312 parents, as “modern” inbred lines, have lost the capacity to profit from AM symbiosis (*e.g.* see Koide et al., 1988; Hetrick et al., 1992), we did observe QTL linked to differences in benefit, implying that it would be possible to select improve the capacity of plants to profit from the symbiosis. Further study will be required to understand the mechanistic basis of the differences we saw in host response. Although the level of AM colonization has been reported to differ among varieties in a number of plant species, it is not necessarily an indicator of greater host response (Kaeppler et al., 2000; Sawers et al., 2017; Watts-Williams et al., 2019; Plouznikoff et al., 2019; Pawlowski et al., 2020). Indeed, the capacity of the host to limit fungal colonization under certain conditions may maximize plant benefit, and it is well established that colonization is reduced under nutrient replete conditions (Karlo et al., 2020; Nouri et al., 2014; Sawers et al., 2017). As much as root-internal fungal growth, the extent of the root-external hyphal network impacts host response (Yao et al., 2001; Munkvold et al., 2004; Schnepf et al., 2008) and may be a factor in benefit variation(Sawers et al., 2017). Recently, it has been shown that certain bacterial taxa exert beneficial and synergistic effects on AM symbiosis, acting as so-called “mycorrhiza helpers” (Frey-Klett et al., 2007; Heijden et al., 2016; Ferreira et al., 2020). Characterization of the root microbiome of *AMF-S* and *AMF-R* families will provide insight into the interplay between AMF, the broader microbial community and host response.

We predicted trade-offs between host dependence and benefit: the genotypes predicted to be the best in AM susceptible families were predicted to be unremarkable or poor in the context of AM resistance, and *vice versa*. At the level of individual loci, examples of antagonistic pleiotropy (Des Marais and Juenger, 2010) were seen for QTL associated with days-to-silking, ear weight, total kernel weight, number of kernels per row and fifty kernel weight. Our identification of intraspecific genetic trade-offs is analogous to patterns of plant nutritional strategy seen at higher taxonomic levels where the differing demands placed on plant anatomy and physiology by direct foraging and engagement in AM symbiosis appear to prevent co-optimization for both (Lambers et al., 2008; Wen et al., 2019). Detailed characterization of root anatomy has suggested that such trade-offs will exist: for example, increasing the proportion of root cortical aerenchyma (root air space) reduces the carbon demand of the root system, promoting foraging efficiency (Postma and Lynch, 2011), but, potentially, at the cost of limiting AMF colonization and host response (Strock et al., 2019). It was somewhat surprising to see a QTL for a morphological trait such as tassel branch number to be conditional on AM susceptibility; such observations may indicate the action of pleiotropic systemic signals. Indeed, the phytohormones so important to the regulation of plant development (including patterning of tassel architecture) are also implicated in the establishment and regulation of AM symbiosis (Gutjahr, 2014; Bonfante and Genre, 2015). For example, disruption of ethylene signalling has been shown to modulate the level of (negative) host response to AMF in *Nicotiana attenuata* (Riedel et al., 2008); a DELLA repressor in the gibberellic acid signalling pathway is essential for AM colonization in rice (Yu et al., 2014); a feedback-loop involving strigolactone modulates the extent of AMF colonization in *Medicago truncatula* (Müller et al., 2019). It may be significant that many of these same signals have been the targets of domestication and plant improvement (Sawers et al., 2018). QTL x AMF effects, including those associated with “trade-off”, were primarily driven by plasticity of the temperate W22 alleles. It was interesting that CML312 alleles, derived from the parent better adapted to the location of our trial, were more stable with respect to the presence or absence of AMF. It has been hypothesised that mutualisms will be strengthened under stressful conditions and that they might play an important role as non-adapted genotypes move into new environments (O’Brien et al., 2018, 2019). Beyond an impact on yield in any given site, the genetic architecture described here implies a role for the AMF in influencing yield stability. In the face of variation and unpredictability in the AM community from site to site, or year to year, “optimization” of the plant host might reasonably place a greater emphasis on consistency over maximum benefit. Although we have found no evidence of a selective pressure to minimize the AM symbiosis in maize breeding, the trade-offs we have described suggest that efforts to optimize plants for better performance either with or without the contribution of microorganisms may not be easily aligned. Ultimately, we may need to make decisions about the types of agroecosystems we want to create and develop our crop varieties accordingly.

## MATERIALS AND METHODS

### Plant material and generation of the F_2:3_ mapping population

An F_2:3_ population was developed from the cross between a W22 stock homozygous for the *hun1-2* allele (*Mutator* insertion mu::1045305 in the gene GRMZM2G099160/Zm00001dwww.maizegdb.org. Ramírez-Flores MR. M.Sc. thesis, 2015) and the subtropical CIMMYT inbred line CML312. The F_1_ was self-pollinated to generate an F_2_ segregating *hun1-2* and W22/CML312 genome content. F_2_ plants were genotyped by PCR to identify homozygous wild-type and mutant individuals, and these self-pollinated to generate susceptible (*AMF-S*) and resistant (*AMF-R*) F_2:3_ families, respectively. F_2:3_ families were increased by sibling-mating. Material was generated in Valle de Banderas, Nayarit, Mexico and Irapuato, Guanajuato, Mexico from 2016 to 2019. Small amounts of seed are available for distribution through direct contact with the authors.

### DNA extraction and PCR genotyping of hun1-2

DNA extraction was performed as described in (Fulton et al., 1995). The genotype at the *Hun* locus was determined by running wild-type and mutant PCR reactions. To amplify the wild-type fragment, a pair of gene-specific primers were used (HUNF01-CGCGAAGAAACGCAGACATTCC and HUNR04-TAACCTGGAGCGAACAGAATCCAC), generating a product of 606 bp. To amplify the mutant fragment a combination of *Mutator* primer and gene-specific primer were used (HUNF03-CTTGGGCGCATTGGAAATTCATCG and RS183-CGCCTCCATTTCGTCGAATCCSCTT), generating a fragment of 839 bp. PCR conditions were: 1 cycle of initial incubation at 94°C for 3 min, 32 cycles of 94°C for 30 sec, 60°C for 30 sec and 72°C for 1 min, and 1 cycle of a final extension at 72°C for 5 min. PCR used the Kapa Taq PCR kit from Kapa Biosystems (Wilmington, Massachusetts, United States) following the manufacturer’s instructions. Products were visualized on 1 % agarose gel.

### Whole genome genotyping and genetic map construction

Approximately 200 F_2_ parents were analyzed using the Illumina MaizeLD BeadChip (https://www.illumina.com/products/by-type/microarray-kits/maize-ld.html). Over 3,000 single nucleotide polymorphisms (SNPs) were detected, with a call rate of> 96% for all samples. The SNP set was processed using TASSEL 5 (Bradbury et al., 2007). SNPs were transformed to ABH format (A: CML312; B: W22) and filtered to remove sites matching any of the following criteria: > 30% missing data; monomorphic sites outside of the *Hun* locus; SNPs outside the *Hun* locus showing segregation distortion (***X***^2^ test, p < 0.05); redundant sites. The markers “5_2269274”, “5_3096229” and “5_4270584” (physical position on chromosome 5 at 2.67, 3.1 and 4.27 Mb, respectively) were used to confirm the expected genotype at the *Hun* locus. The genetic map was built in R (R core team 2020) using R/ASMAP (Taylor and Butler, 2017) and R/qtl (Broman et al., 2003); (Broman and Sen, 2009a).

### Field Evaluation and phenotypic analysis

The F_2:3_ population was evaluated in the summer of 2019 at the UNISEM experimental station in Ameca, Jalisco, Mexico (20.78, −105.243). Three complete blocks of 73 *AMF-S* and 64 *AMF-R* families were evaluated. Within each block, *AMF-S* and *AMF-R* families were alternated. The order of the families within the *AMF-S* or *AMF-R* sub-populations was randomized within each block. For each replicate, 45 seeds of each family were sown in a plot of three 2 m long rows. Only the second row of each plot was considered for evaluation, the first and third rows providing a buffer between adjacent families. Phenotypic data were collected at flowering and after harvest as follows: stand count (STD), number of plants per row; days to anthesis (DTA), number of days from planting until anthers visible on the main spike of half of the plants in the row; days to silking (DTS), number of days from planting until silks visible on half of the plants in the row; anthesis-silking interval (ASI), difference in days between anthesis and silking; plant height (PH), distance in cm from the ground to the flag leaf; tassel branch number (TBN), number of primary lateral branches originating from the main spike; ear weight (EW), the weight in g of the ear; ear length (EL), distance in centimeters from the base to the tip of the ear; ear diameter (ED), the diameter in cm at the middle part of the ear; kernel row number (KRN), number of rows in the middle part of the ear; kernels per row (KPR), number of kernels in a single row from the base to the tip of the ear; cob diameter (CD), diameter in cm at the middle part of the cob; fifty kernel weight (FKW), weight in g of 50 randomly selected kernels; total kernel weight (TKW), weight in g of all kernels on the ear; total kernel number (TKN), estimated from TKW/FKW * 50.

### Image analysis

Image-based phenotyping method was used to quantify the size and shape of maize ear, cob, and kernel. Ears and cobs (1-3) from each plot, and approximately 50 kernels from associated ears, were scanned using an Epson Perfection V600 scanner at 1200 dots per inch. Scanned images were uploaded to Cyverse (Merchant et al., 2016) and analyzed using an automatic pipeline to quantify ear, cob, and kernel attributes (Miller et al., 2017). Width profiles of ear and cob and contour data of kernels were extracted in R to compute their shapes.

### Principal Component Analysis

Principal Component Analysis (PCA) was conducted to analyze major sources of plant phenotypic variance and to visualize the relationships among all measured phenotypic traits across two maize families. PCA was performed using the prcomp function in R.

### QTL mapping and whole genome prediction

QTL mapping was performed using the R/qtl package (Broman et al., 2003) on a population composed of 73 AMF-S and 64 AMF-R F2:3 families. Single-QTL standard interval mapping was run using the scanone function with Haley-Knott regression, with the genotype at *Hun* (wt =1; *Hun* = 0) treated as a covariate (hereafter, AMF). To assess QTL x AMF interaction (Broman and Sen, 2009), QTL analysis was performed under four different models: separate analyses for AMF-S (H_0wt_) and AMF-R (H_0hun_) subpopulations; considering AMF as an additive covariate (H_a_) and considering AMF as an interactive covariate (H_f_). The LOD significance threshold for each trait and model was established with a permutation test (α = 0.05, 1000 permutations). Evidence of QTL x AMF interaction was obtained by comparing H_f_ and H_a_ models, where the difference LOD_i_ = LOD_f_ - LOD_a_ was considered with reference to the threshold difference LOD_thr_i_ = LOD_thr_f_ - LOD_thr_a_ for evidence of possible interaction. Individual QTL were combined into multiple-QTL models on a per trait basis. Where evidence was found of QTL x AMF interaction in the single scan, the interaction was also included in the multiple-QTL model. Multiple-QTL models were evaluated using the fitqtl function and non-significant (p < 0.1) terms removed according to the drop-one table.

To compare the potential of a given genotype in the presence or absence of AM colonization, we estimated the performance of the 137 families for both levels of AMF using two approaches. In the first approach, we generated fitted values from the multiple-QTL models, under both levels of the AMF covariate. In the second approach, we performed whole genome prediction using the R/rrBLUP package (Endelman, 2011). Missing genotypes were imputed with the A.mat function and marker effects and BLUE values were obtained with the mixed.solve function. The observed values for AMF-S families were used to train a mycorrhizal model, that was then applied to the genotypes of the AMF-R families; the observed values from the AMF-R families were used to train a non-mycorrhizal model, that was then applied to the genotypes of the AMF-S families. For interpretation of the results, predicted values were compared for all genotypes in both levels of AMF, avoiding possible bias in extreme observed values.

## ACKNOWLEDGEMENTS

We thank Beda Angehrn and Mario Rivera (UNISEM) for assistance with field evaluation, Cruz Robledo (PV Winter Nurseries) for support with generating material, and Jessica Carcaño-Macias for technical support and seed stock management. We acknowledge Karina Picazarri-Delgado for assistance with preliminary characterization of the *hun* mutants and Nathan Miller for his help with image analysis. This study was funded by the Mexican National Commission for the Study and Use of Biodiversity (CONABIO) project *Impact of native arbuscular mycorrhizal fungi on maize performance* (N° 62, 2016-2018). M. Rosario Ramírez-Flores was supported by a Ph.D. scholarship from the Mexican National Council of Science and Technology (CONACYT).

