## Supplemental_Information for "The genetic architecture of host response reveals the importance of arbuscular mycorrhizae to maize cultivation"

### Supplementary Figures

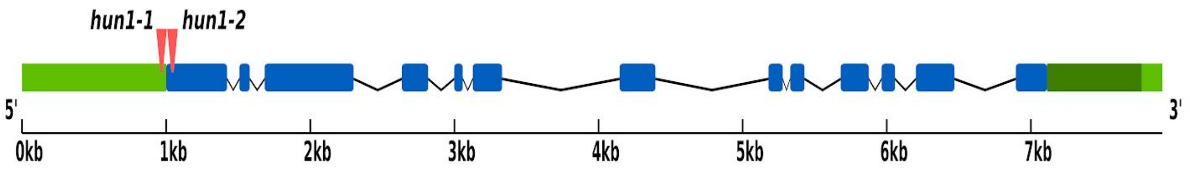


**Figure S1. Location of *Mutator* insertions in *Hun1***. Representation of the gene GRMZM2G099160/Zm00001d012863 (B73 v4 genome). The transcript shown is GRMZM2G099160_T01.1, this version has 13 coding exons, transcription length is 3422 bps and translation length of 926 residues. Two *Mutator* insertions were identified in the maize *Hun1* gene from the Uniform_Mu collection (www.maizegdb.org; McCarty *et al.*, 2005; Ramírez-Flores, 2015). Sequence flanking the event *mu1018108* (stock UFMu-01071) was located in chromosome 5 at 1298802..1298810 bp (B73 v4 genome), 44 bp upstream of the *Hun1* translational start site, and was designated *hun1-1*. Sequence flanking the event *mu1045305* (stock UFMu-05472) was located in chromosome 5 at 1298719...1298727 bp, 39 bp downstream of the translational start site, and was designated *hun1-2*. Red triangles represent *Mutator* insertions, blue boxes correspond to exons, black lines the introns, light green boxes flanking regions in HUN and dark green box to the annotated 3'UTR.

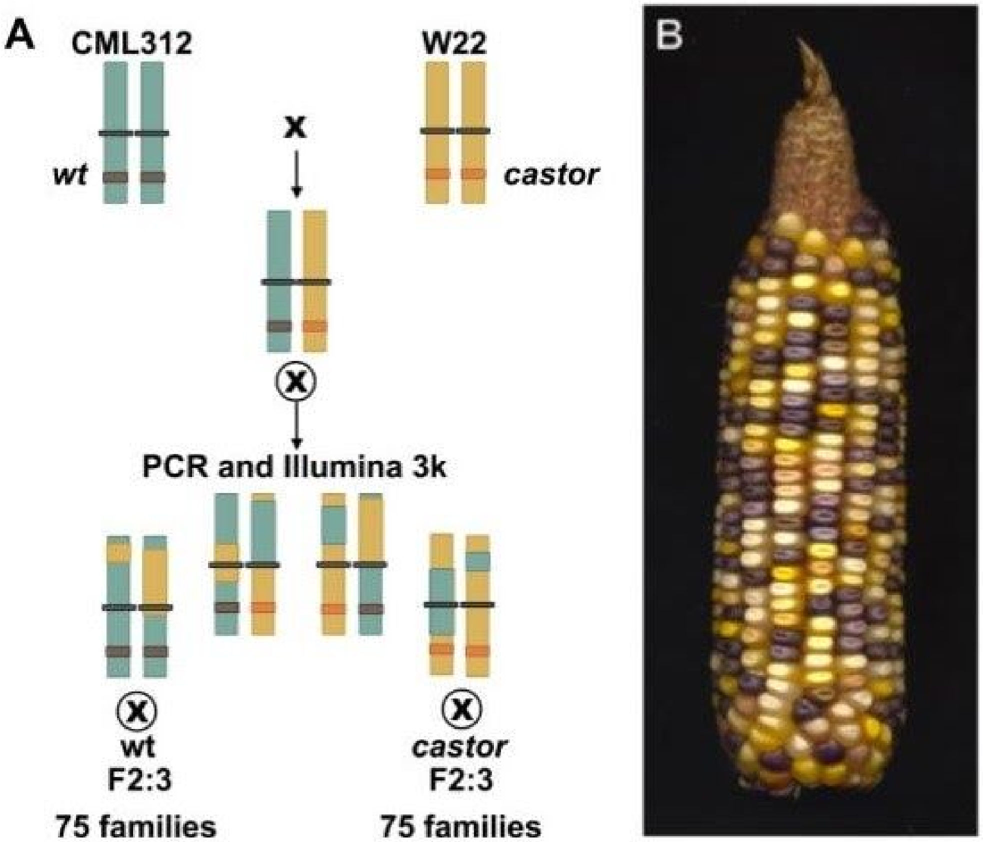


**Figure S2. Generation of the $F_{2:3}$ population.** A) $F_1$ was generated from the cross between the CML312 inbred line and a plant homozygous for the *hun1-2* allele in the W22 background. The $F_1$ was self-pollinated to generate a stock segregating *hun1-2* along with CML312 and W22 genome content. Homozygous wild-type and mutant individuals were identified by PCR genotyping and self-pollinated to generate ~ 75 $F_{2:3}$ families.. Outside of the *Hun1* locus, resistant and susceptible families were anticipated to segregate equally for W22 and CML312 genome content, and thereby provide a phenotypic range against which to assess mycorrhiza response. Additional tissue from the $F_2$ parents was used for genotypic analysis on an Illumina 3047 SNP microarray chip (www.illumina.com/products/by-type/microarray-kits/maize-ld.html) and a genetic linkage map was constructed for downstream QTL analysis. Aside from the expected segregation distortion at the *Hun1* locus, the genetic markers showed Mendelian segregation across families. Seed of all 137 $F_{2:3}$ families was increased by sibling mating for use in field evaluation. B) An $F_2$ ear from the cross of the color-converted *bz-mum9* Uniform Mu stock carrying *hun1-2* and the white-kernelled subtropical inbred line CML312.

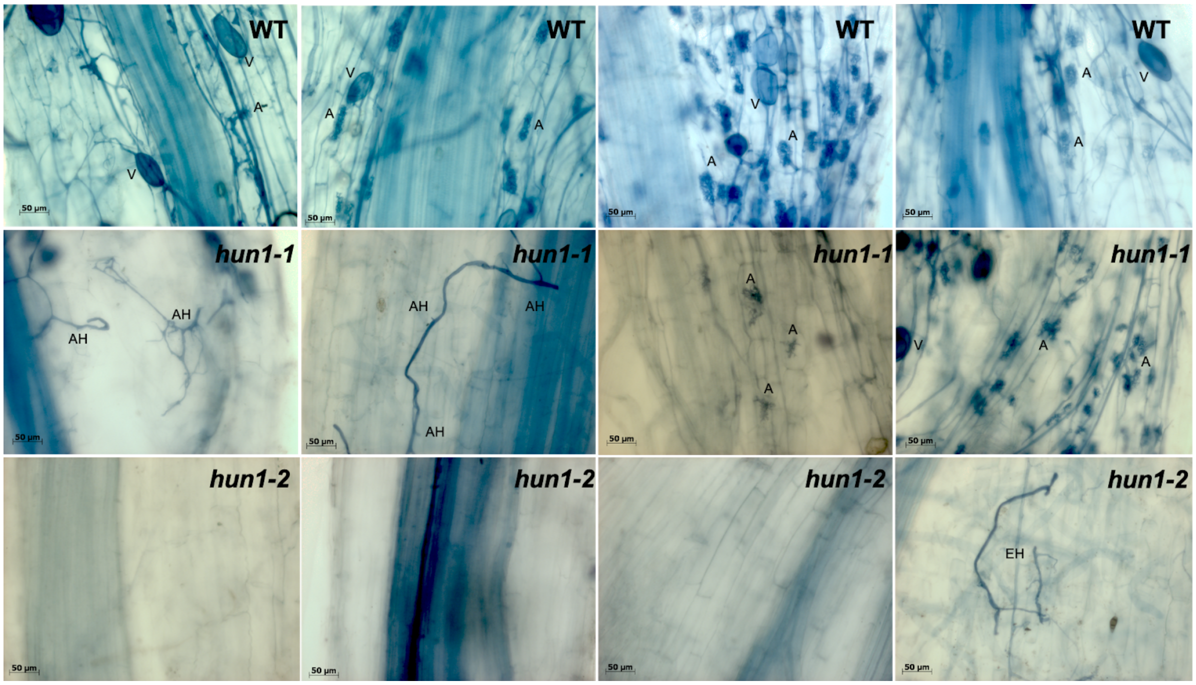


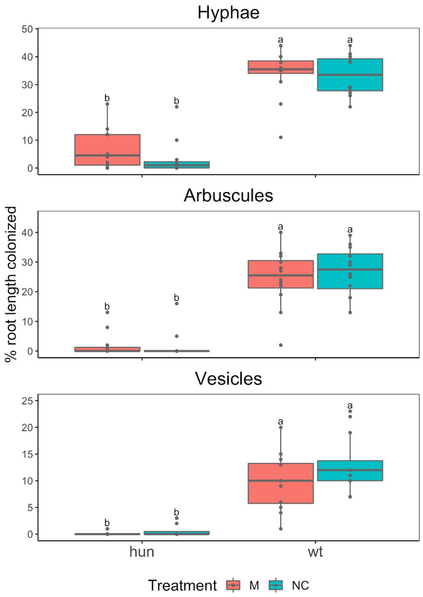


**Figure S3. *hun1* mutants are resistant to colonization by AMF. Above,** Greenhouse characterization of wild-type (WT) and *hun* mutant roots, harvested at 56 DAE and colonized with *Rhizophagus irregularis*. (A) to (D) Roots of the WT genotypes B73 (A), W22 (B), WT1-1 (C) and WT1-2 (D), show colonization by hyphae, vesicles and arbuscules. (E) to (H) Roots of *hun1-1*, (E) and (F) show colonization by hyphae and abortive hyphae are observed, (G) and (H) colonization by hyphae, vesicles and arbuscules are observed. (I) to (L) Roots of *hun1-2,* extraradical hyphae are observed and structures such as vesicles and arbuscules are not observed. A = arbuscules, V = vesicles, AH = abortive hyphae, EH = extraradial hyphae. **Left,** Field evaluation of *hun1-2* mutants, summer of 201, Irapuato, Mexico. Two complete blocks with 4 treatments was used: 1) AMF-S families without inoculation of AMF, 2) AMF-R families without inoculation of AMF, 3)AMF-S families with inoculation of AMF and 4) AMF-R families with inoculation of AMF. The AMF used was an inoculum from the commercial consortium BioMic which includes: *Glomus constrictum*, *Glomus geosporum*, *Glomus tortuosum*, *Acaulospora scrobiculata*, *Gigaspora margarita* and *Glomus sp*.

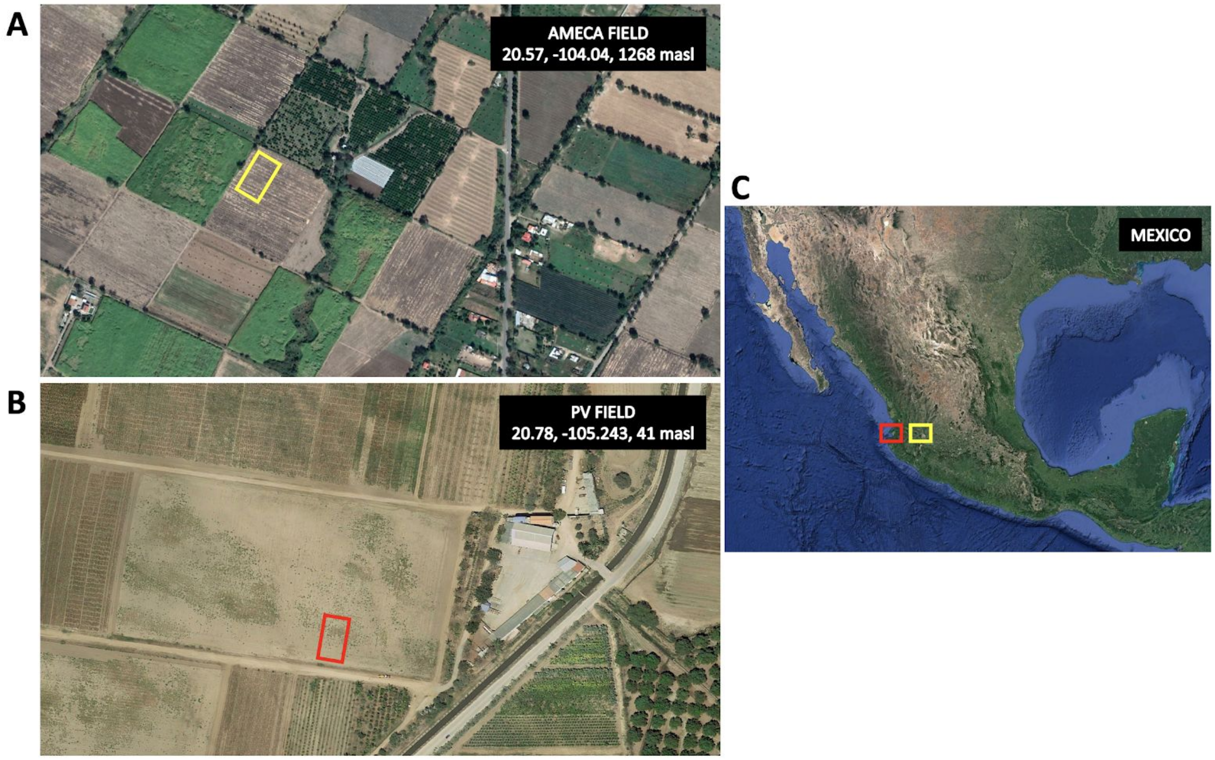


**Figure S4. Location of experimental fields**. A) Satellite view of the Ameca site used for the principal field evaluation. Ameca is located in Jalisco state in the central-western of México. Yellow rectangle represents the area of the experiment. The latitude of the site is 20.57, longitud is -104.04 and elevation 1268 meters above the sea level (B) Satellite view of the Puerto Vallarta (PV) site used for population generation and preliminary high-input evaluation. The PV site is located in Nayarit state on the Pacific Coast. The latitude of the site is 20.78, longitud is -105.243 and elevation 41 meters above sea level. Red rectangle shows the field area (C) Mexico showing the two sites.

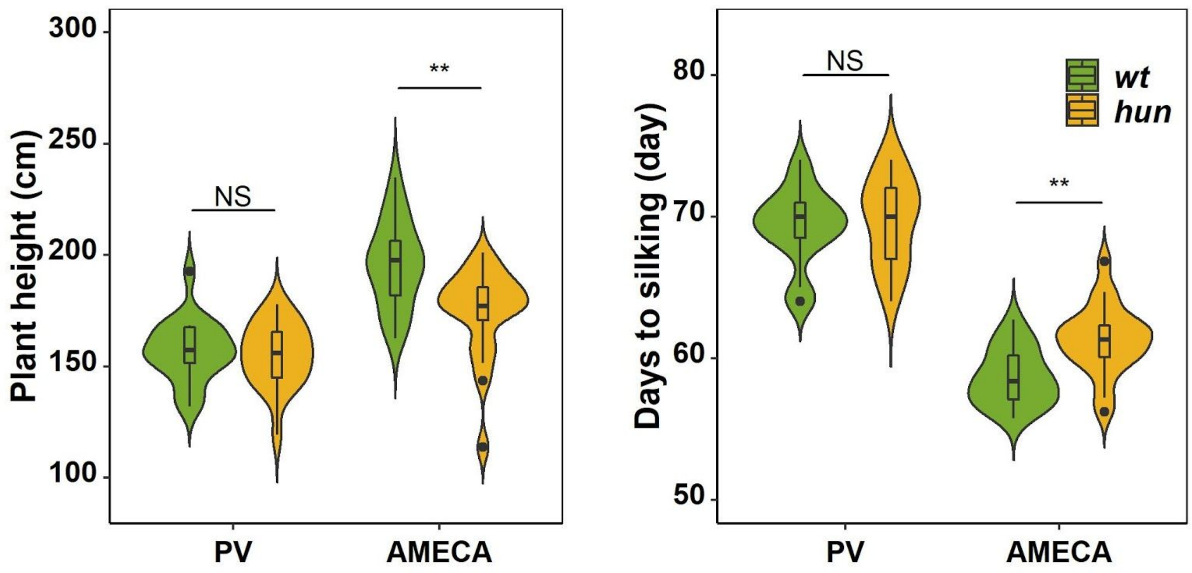


**Figure S5**. Evaluation of *AMF-S* (wt) and *AMF-R* (hun) families in heavily managed (PV) and a rain-fed, medium input (Ameca) fields. Prior to full evaluation a subset of 20 *AMF-S* and 20 *AMF-R* families was evaluated in a winter season in a heavily managed (fertigation, herbicide, pesticide) field site near Puerto Vallarta on the Mexican Pacific Coast. No difference was seen between *AMF-S* and *AMF-R* families in plant height or days-to-silking. There were clear environmental differences with the Ameca site used for the full evaluation where, importantly, *AMF-S* and *AMF-R* families were distinct. The box in violin plots represents the interquartile range with the horizontal line representing the median and whiskers representing 1.5 times the interquartile ranges. The shape of the violin plot represents probability density of data at different values along the y-axis. Results were based on two-group Wilcoxon tests with Bonferonni adjusted P-values. Note: *: $p < 0.05$; **: $p < 0.01$; ***: $p < 0.001$; NS: not significant.

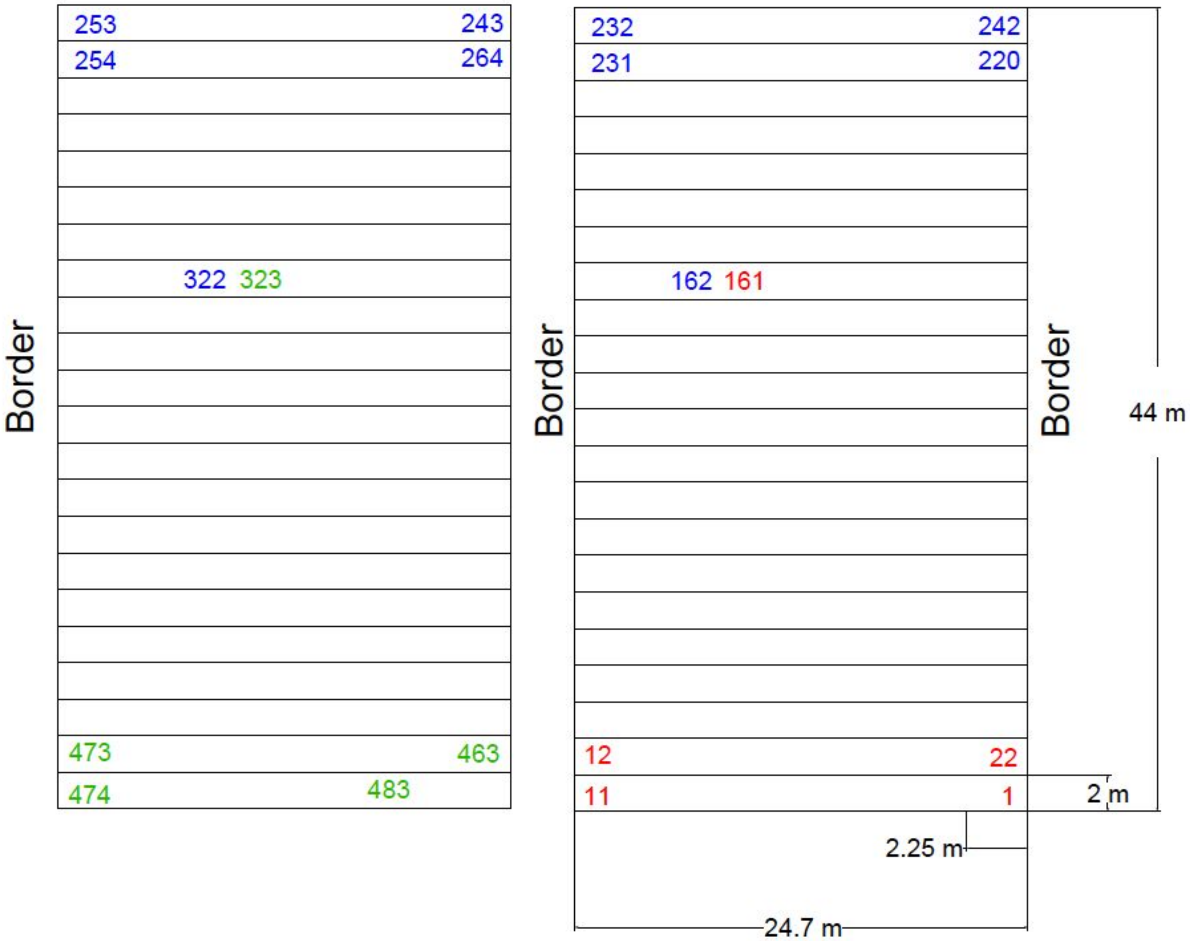


**Figure S6. Plan of the Ameca field design.** Summer experimental field 2019, located in Ameca, Jalisco. Three complete blocks were evaluated. Forty-five seeds were sown from per 3 row plot. AMF-S and AMF-R families were alternated with the order of the families within each subpopulation randomized within each block. The rows marked with red, blue and green correspond to blocks 1, 2 and 3, respectively. A commercial UNISEM hybrid was planted on the border of the experiment and used as a check throughout the field. The field was fertilized at planting with 250 kg / Ha of diammonium phosphate (DAP) as a source of nitrogen and phosphorus (18-46-00, NPK). A further application of 250 kg / Ha of urea was given (46-00-00, NPK) at 40 days after planting.

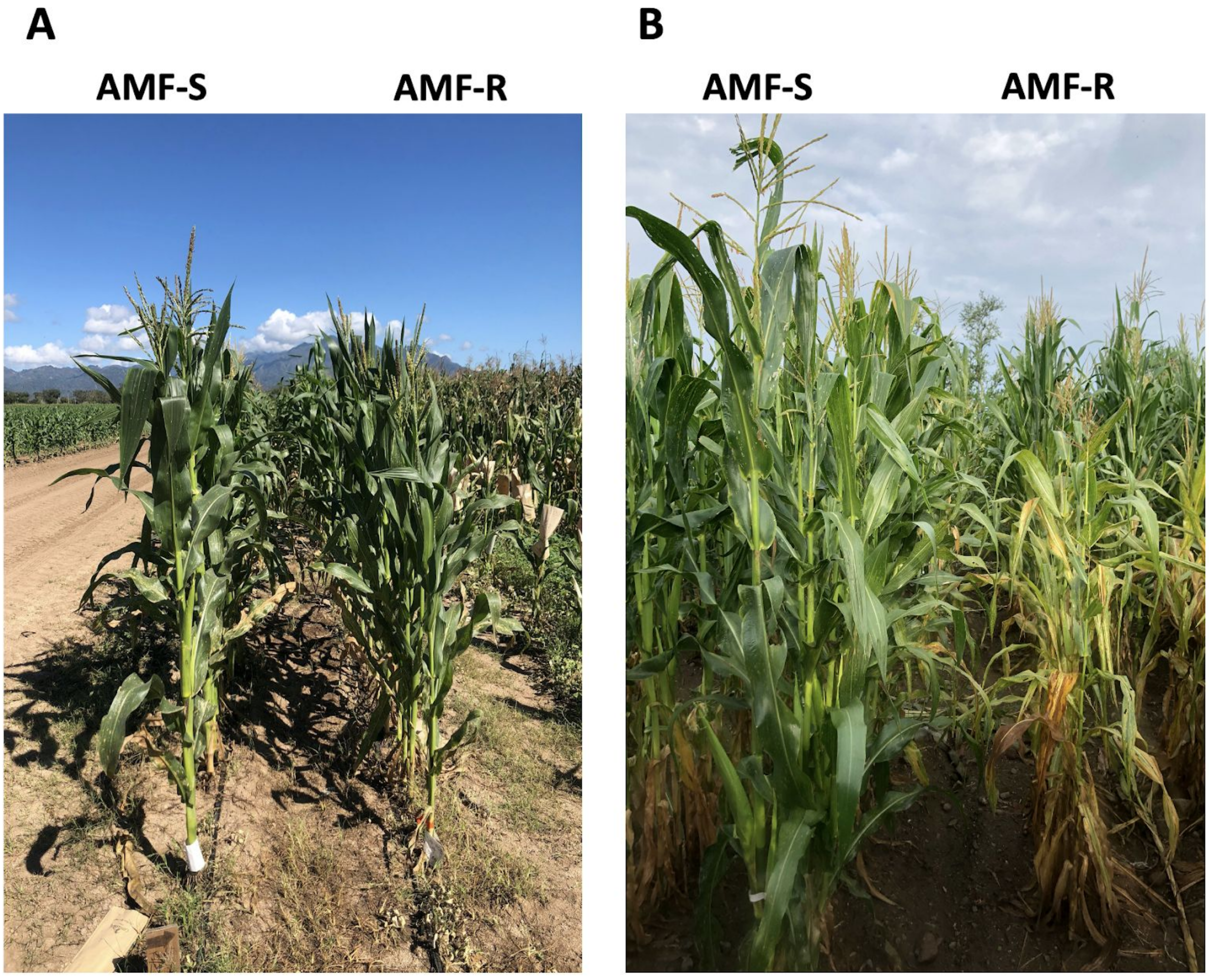


**Figure S7. Overview of representative AMF-S and AMF-R families in high- and medium- input sites.** A) Winter PV field in 2020, located in Nayarit, Mexico. B) Summer Ameca field in 2019, located in Jalisco, Mexico.

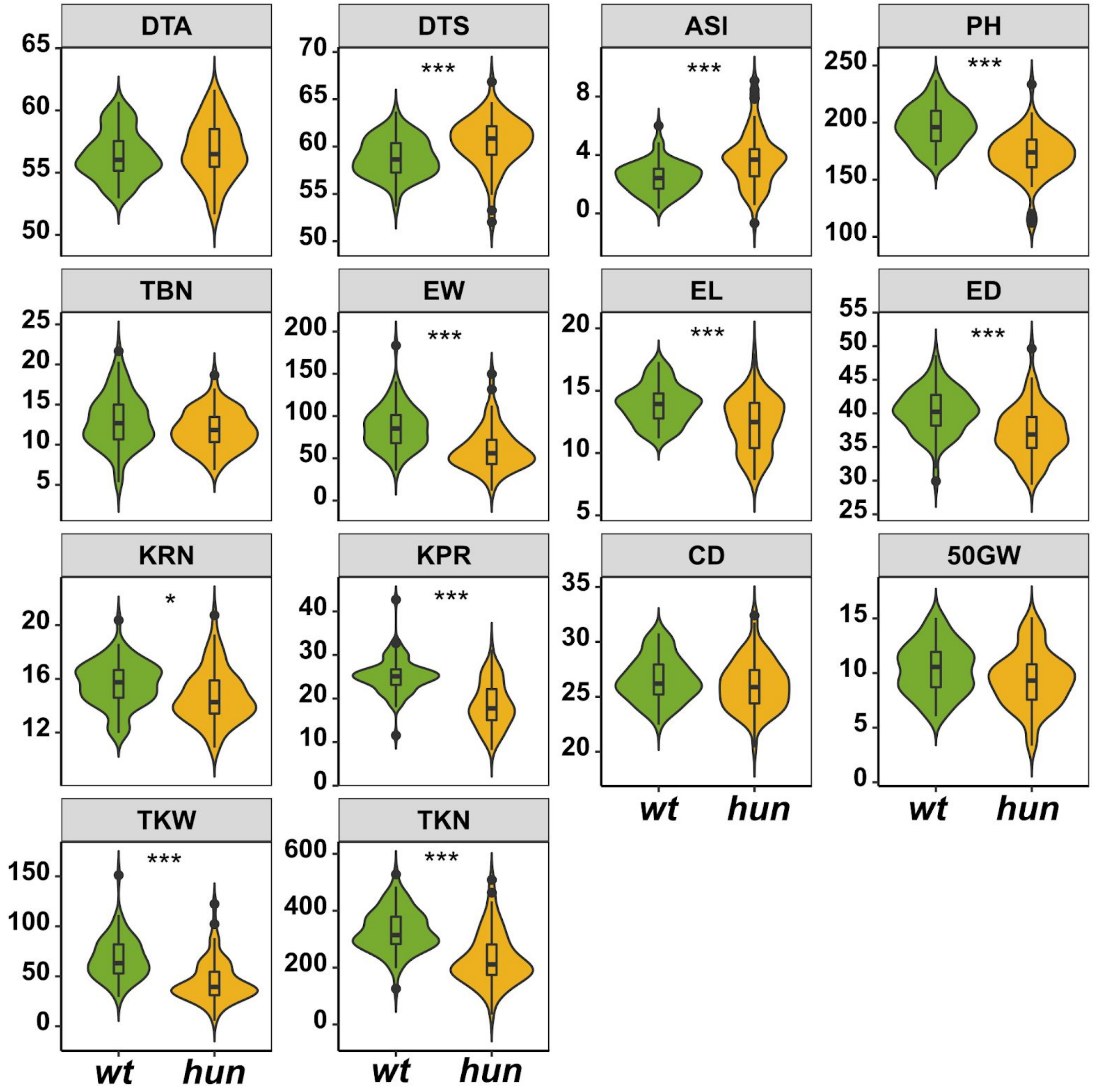


**Figure S8.** Comparison of means of plant phenotypic traits (as shown in Table X.) between AMF susceptible (wt) and resistant (hun) families. The box in violin plots represents the interquartile range with the horizontal line representing the median and whiskers representing 1.5 times the interquartile ranges. The shape of the violin plot represents probability density of data at different values along the y-axis. Results were based on two-group Wilcoxon tests with Bonferonni adjusted P-values. Note: *: $p < 0.05$; **: $p < 0.01$; ***: $p < 0.001$; NS: not significant.

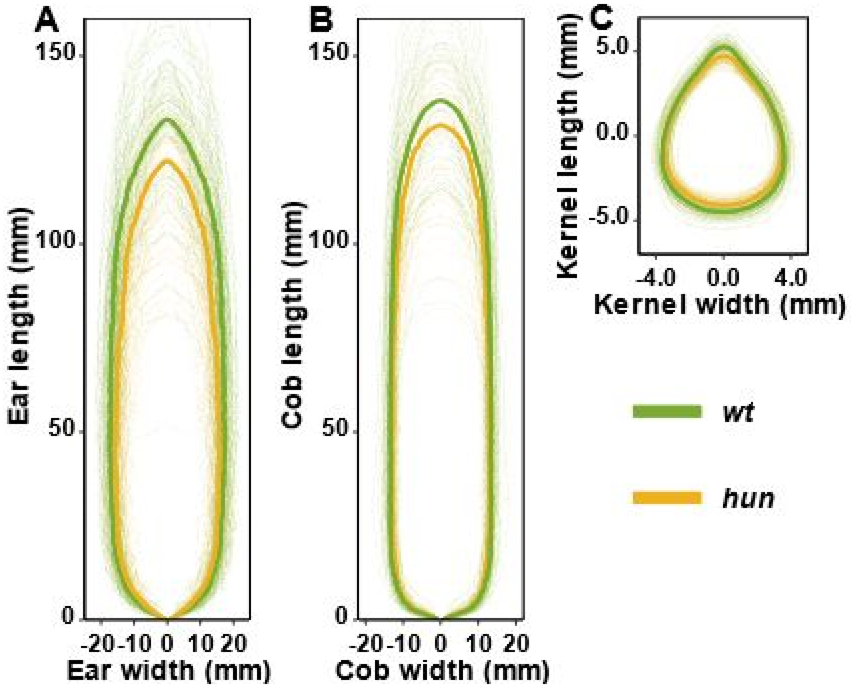


**Figure S9. Ear and kernel image analysis.** The shape of maize ear (A), cob (B), and kernel (C) of AMF susceptible (wt) and resistant (hun) families. Median shapes of ear, cob, and kernel from two families were shown in thick green and yellow lines, whereas individual genotypes of two groups were shown in thin, semi-transparent green and yellow lines.

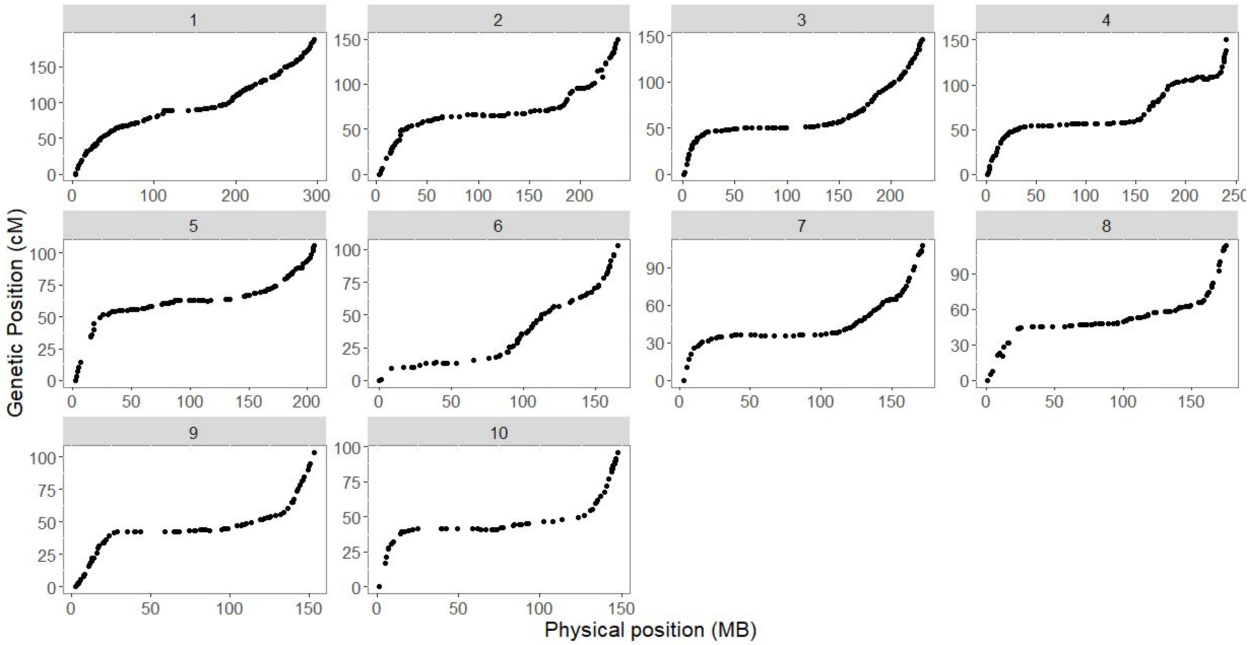


**Figure S10. Genotypic analysis.** Plot of genetic (y axis) and physical (x axis) position for 1050 genetic markers that compose the genetic map of the CML312 x W22 *hun* F2 population.

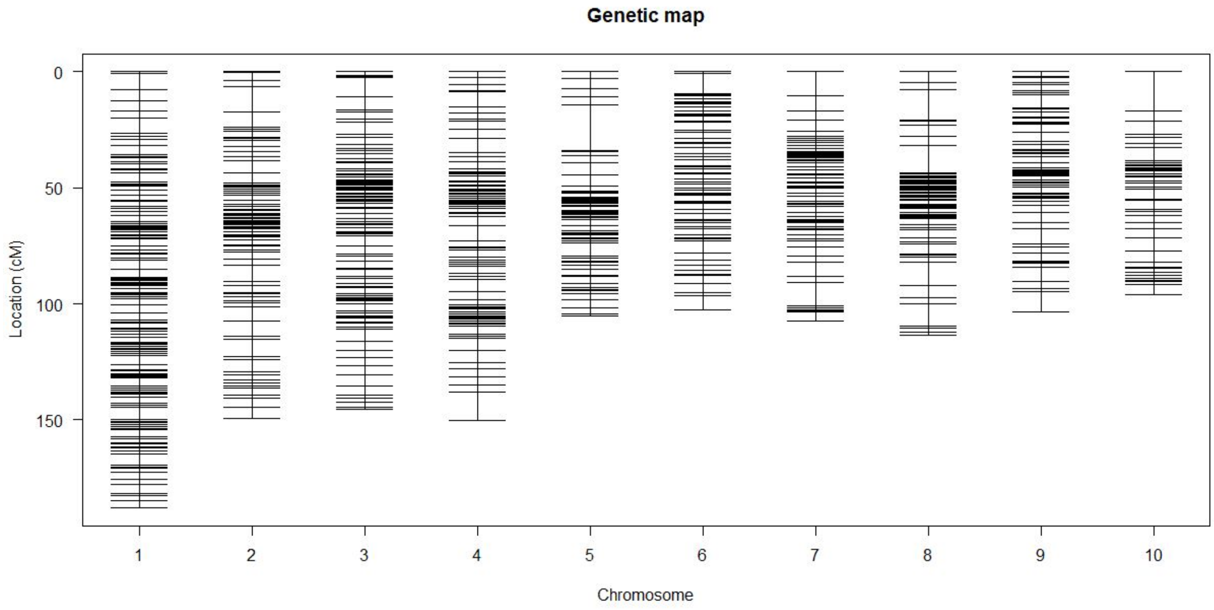


**Figure S11. Genetic map.** Plot representing the estimated genetic map and the marker distribution across the ten chromosomes of maize for the CML312 x W22 *hun* F2 population.

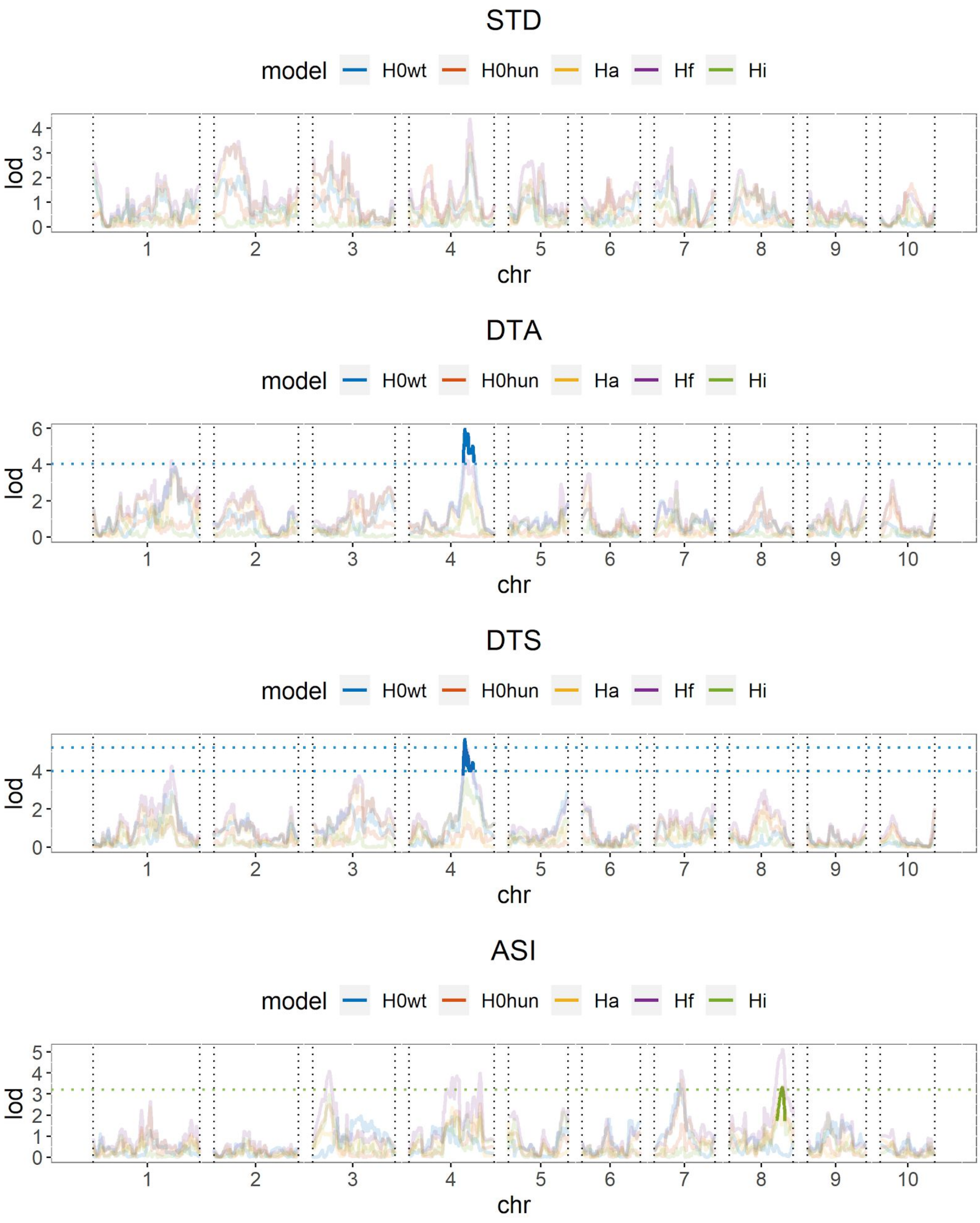
STD
model
H0wt
H0hun
Ha
Hf
Hi
lod
chr
DTA
DTS
ASI

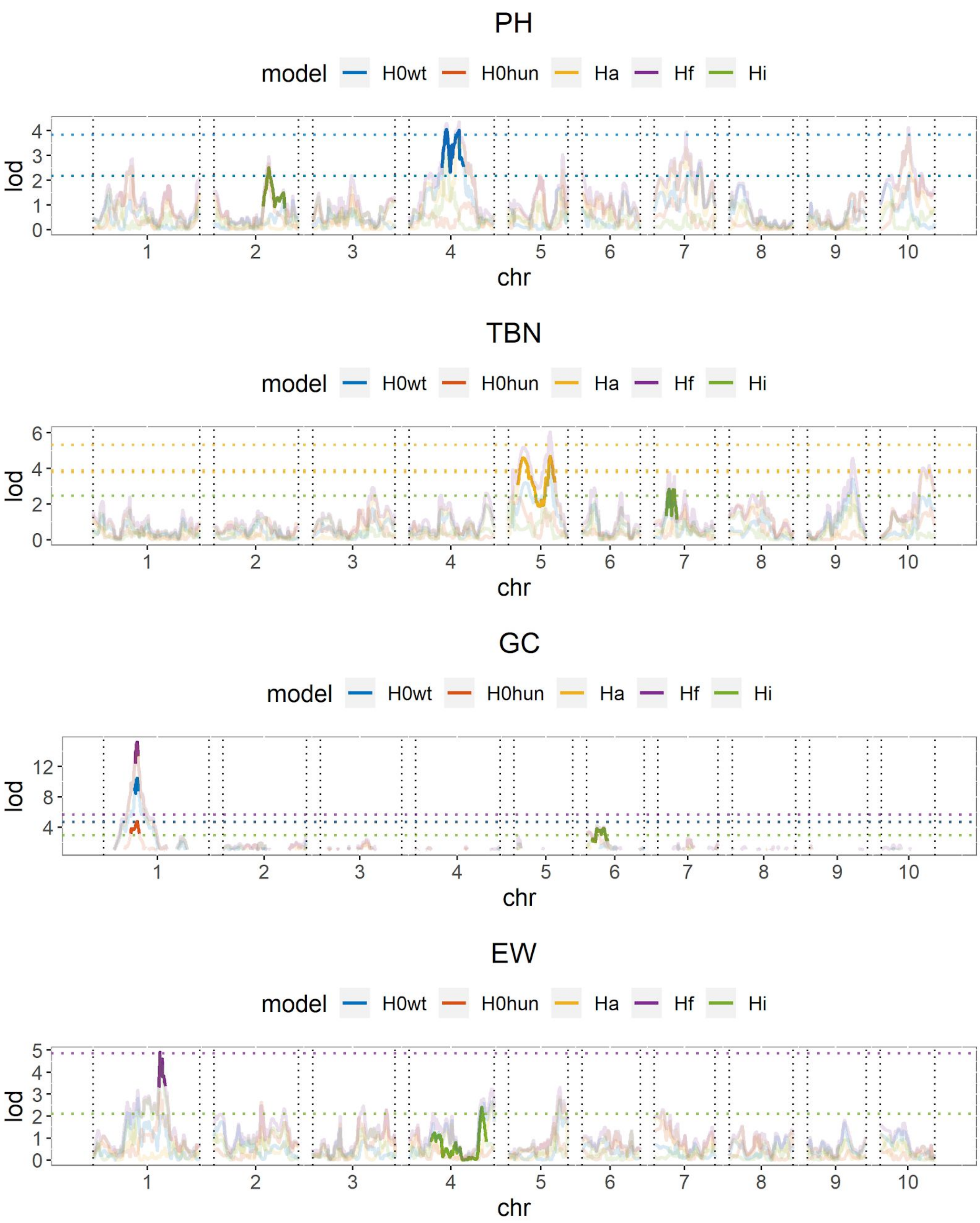
PH
model H0wt H0hun Ha Hf Hi
lod
0 1 2 3 4
chr
1 2 3 4 5 6 7 8 9 10
TBN
model H0wt H0hun Ha Hf Hi
lod
0 2 4 6
chr
1 2 3 4 5 6 7 8 9 10
GC
model H0wt H0hun Ha Hf Hi
lod
4 8 12
chr
1 2 3 4 5 6 7 8 9 10
EW
model H0wt H0hun Ha Hf Hi
lod
0 1 2 3 4 5
chr
1 2 3 4 5 6 7 8 9 10

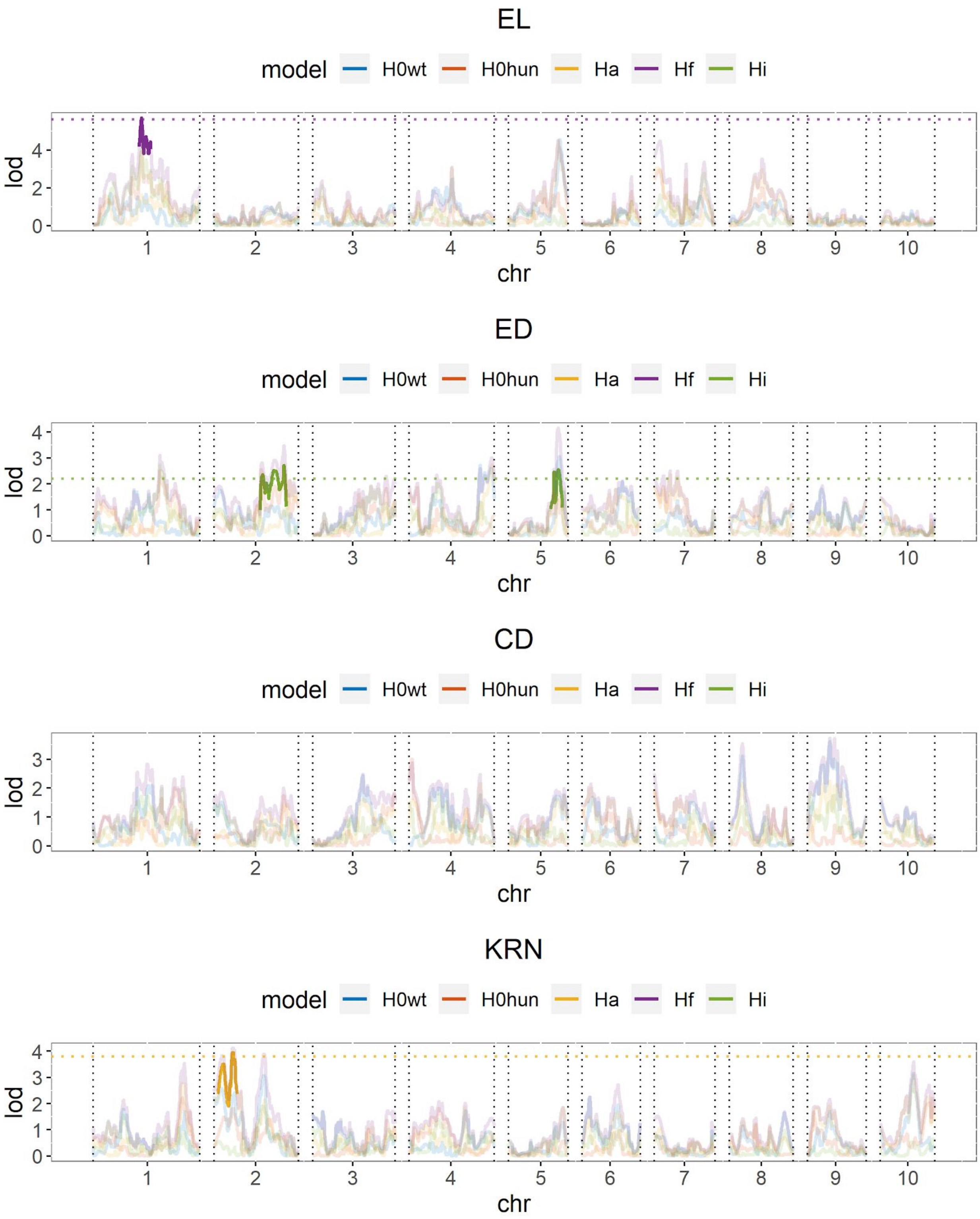

EL
model
H0wt
H0hun
Ha
Hf
Hi
lod
0
2
4
1
2
3
4
5
6
7
8
9
10
chr
ED
model
H0wt
H0hun
Ha
Hf
Hi
lod
0
1
2
3
4
1
2
3
4
5
6
7
8
9
10
chr
CD
model
H0wt
H0hun
Ha
Hf
Hi
lod
0
1
2
3
1
2
3
4
5
6
7
8
9
10
chr
KRN
model
H0wt
H0hun
Ha
Hf
Hi
lod
0
1
2
3
4
1
2
3
4
5
6
7
8
9
10
chr

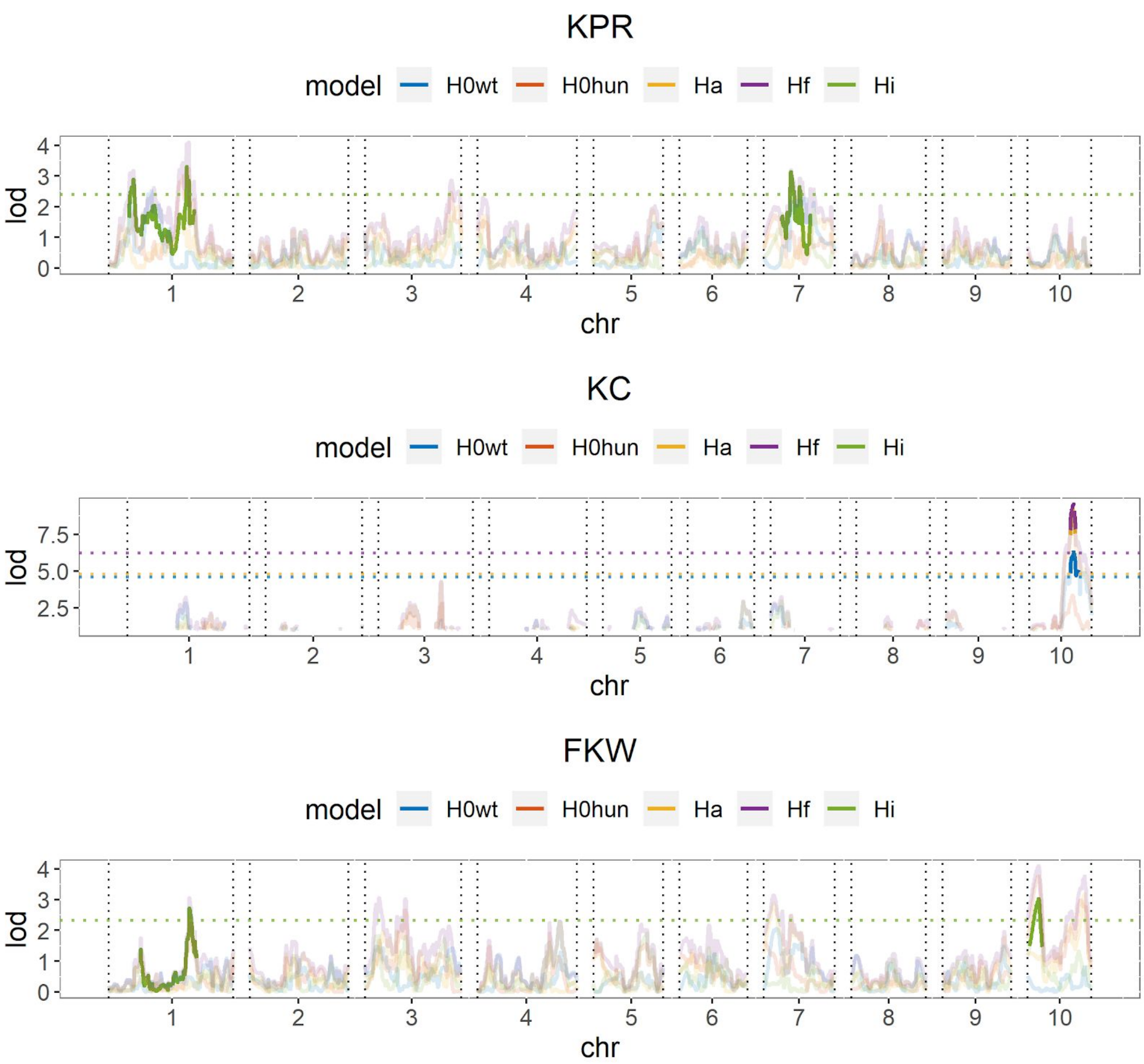
KPR
model H0wt H0hun Ha Hf Hi
lod
chr
KC
model H0wt H0hun Ha Hf Hi
lod
chr
FKW
model H0wt H0hun Ha Hf Hi
lod
chr

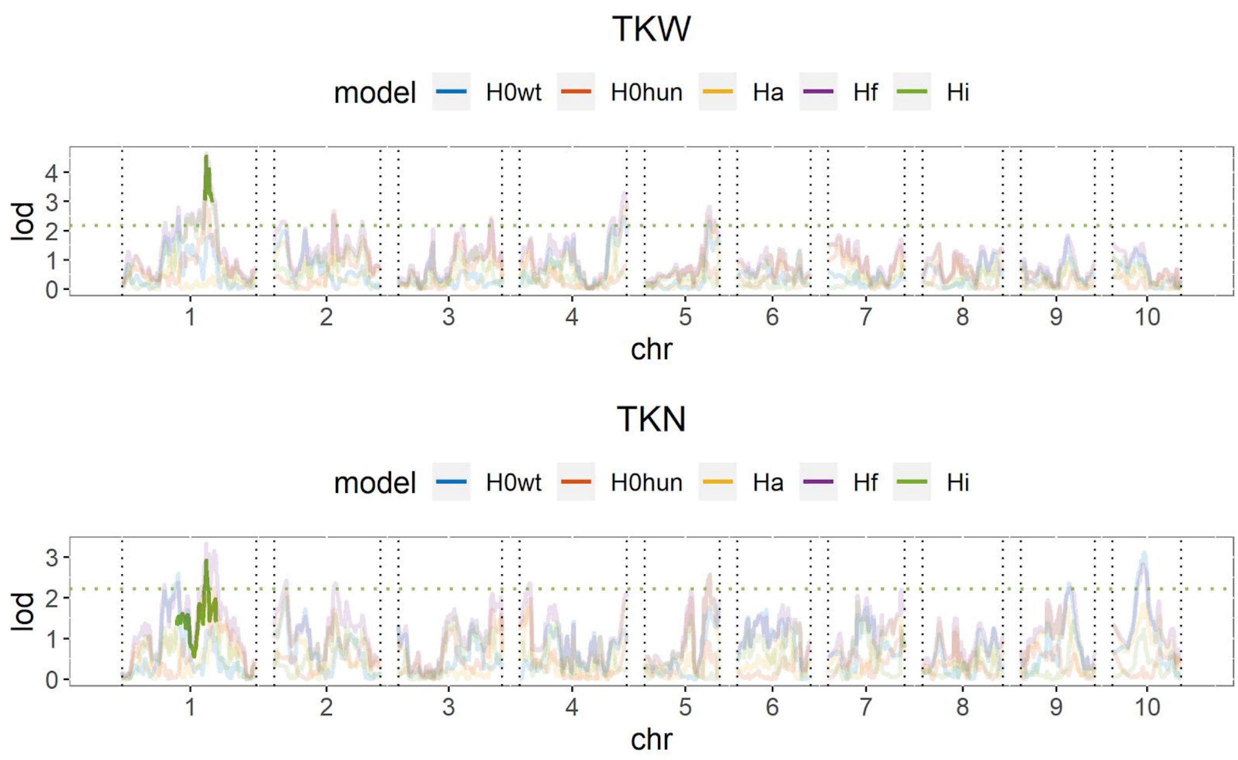


**Figure S12: QTL analysis.** LOD plots of the single-scan QTL performed on the 17 traits studied (see Table S2 for trait codes). The color of the line represent the different models considered: AMF-S families only ($H_{0wt}$, blue line), AMF-R only ($H_{0hun}$, red line), with AMF as an additive covariate ($H_a$, yellow line) and as a interactive covariate ($H_f$, purple line), and the evidence of interaction ($H_i$, green line). The solid line represents the QTL and its 1-LOD drop confidence interval. The horizontal dotted line represents the LOD threshold obtained with a 1000 permutation test ($\alpha = 0.05$) for $H_{0wt}$, $H_{0hun}$, $H_a$ and $H_f$ and calculated as $LOD\_thr_i = LOD\_thr_f - LOD\_thr_a$ for $H_i$.

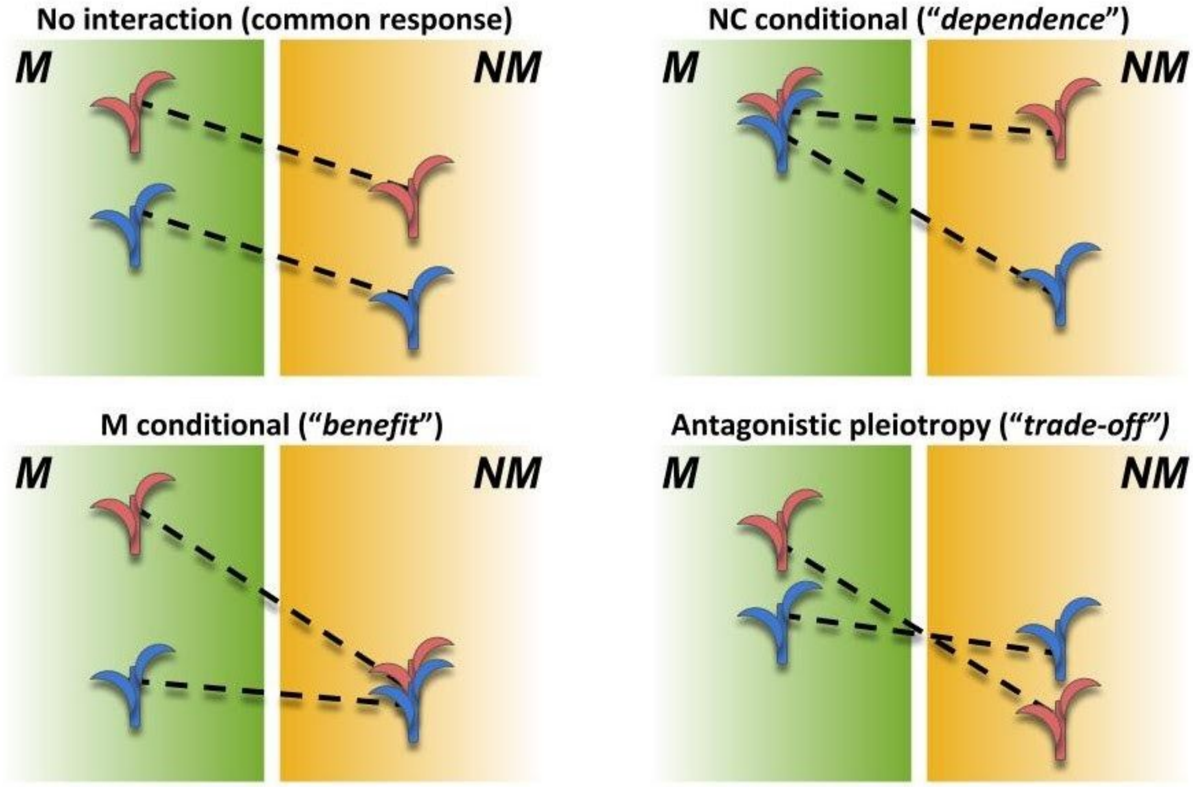


**Figure S13. Scenarios in AMF x genotype interaction.** Theoretical performance (arbitrary units; y axis) of mycorrhizal (M) and non-mycorrhizal (NM) plants of two genotypes (red and blue). The difference between M and NM defines response for a given genotype. Top left) No AMF x genotype interaction. Both genotypes respond equally to AMF; the difference between varieties is constant between M and NM. Top right) NM conditional. Response is higher in the blue genotype; difference between genotypes is conditional on NM growth; response variation is driven by greater dependence of the blue genotype. Bottom left) M conditional. Response is higher in the red genotype ; difference between genotypes is conditional on M growth; response variation is driven by greater benefit of the blue genotype. Bottom right) Antagonistic pleiotropy. Response is higher in the red genotype; difference between the genotypes is expressed in both M and NM, but the sign of the effect changes; the red genotype is superior in M growth but inferior in NM growth.

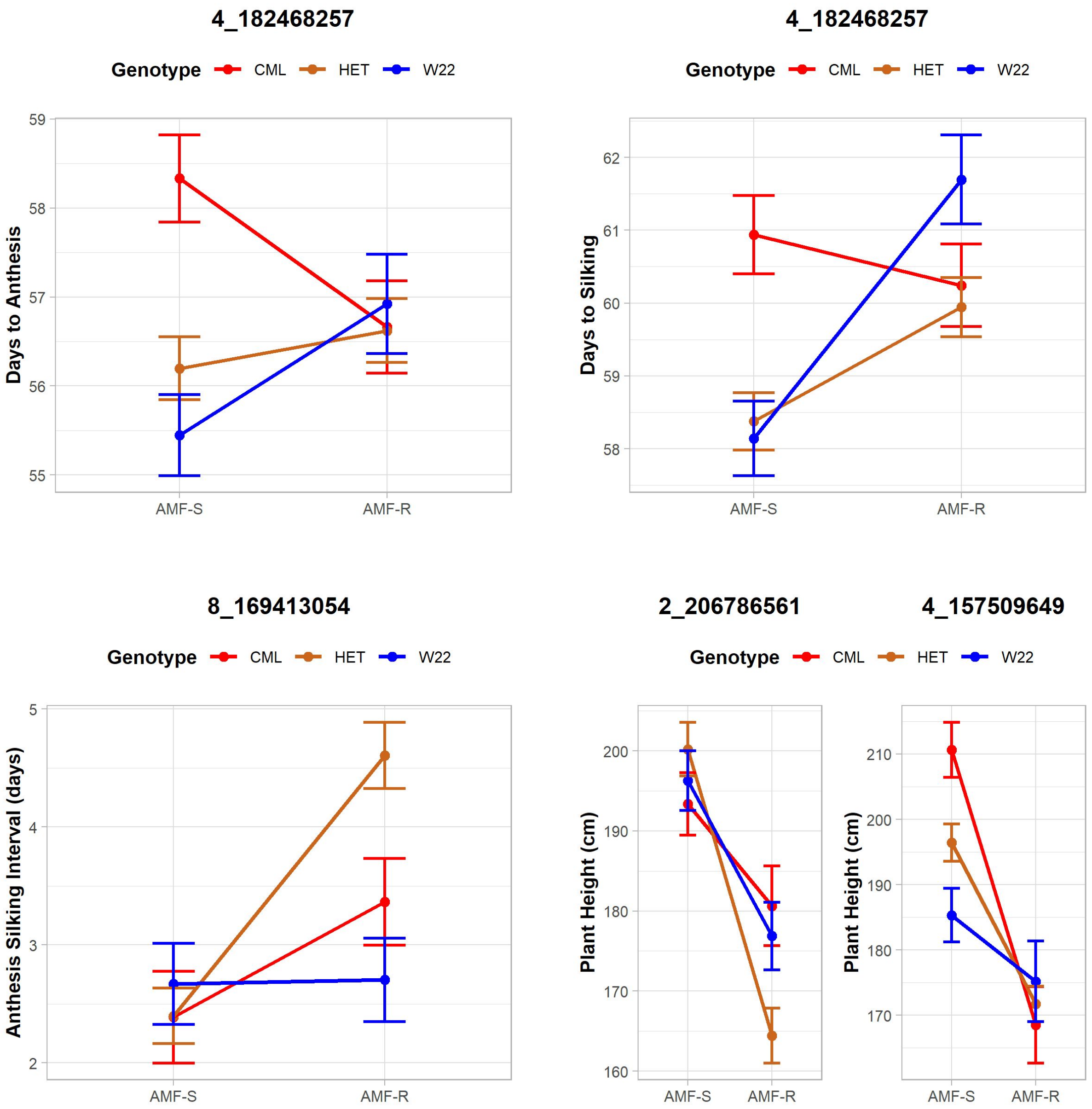
4_182468257
Genotype CML HET W22
Days to Anthesis
59
58
57
56
55
AMF-S
AMF-R
4_182468257
Genotype CML HET W22
Days to Silking
62
61
60
59
58
AMF-S
AMF-R
8_169413054
Genotype CML HET W22
Anthesis Silking Interval (days)
5
4
3
2
AMF-S
AMF-R
2_206786561
4_157509649
Genotype CML HET W22
Plant Height (cm)
200
190
180
170
160
AMF-S
AMF-R
Plant Height (cm)
210
200
190
180
170
AMF-S
AMF-R

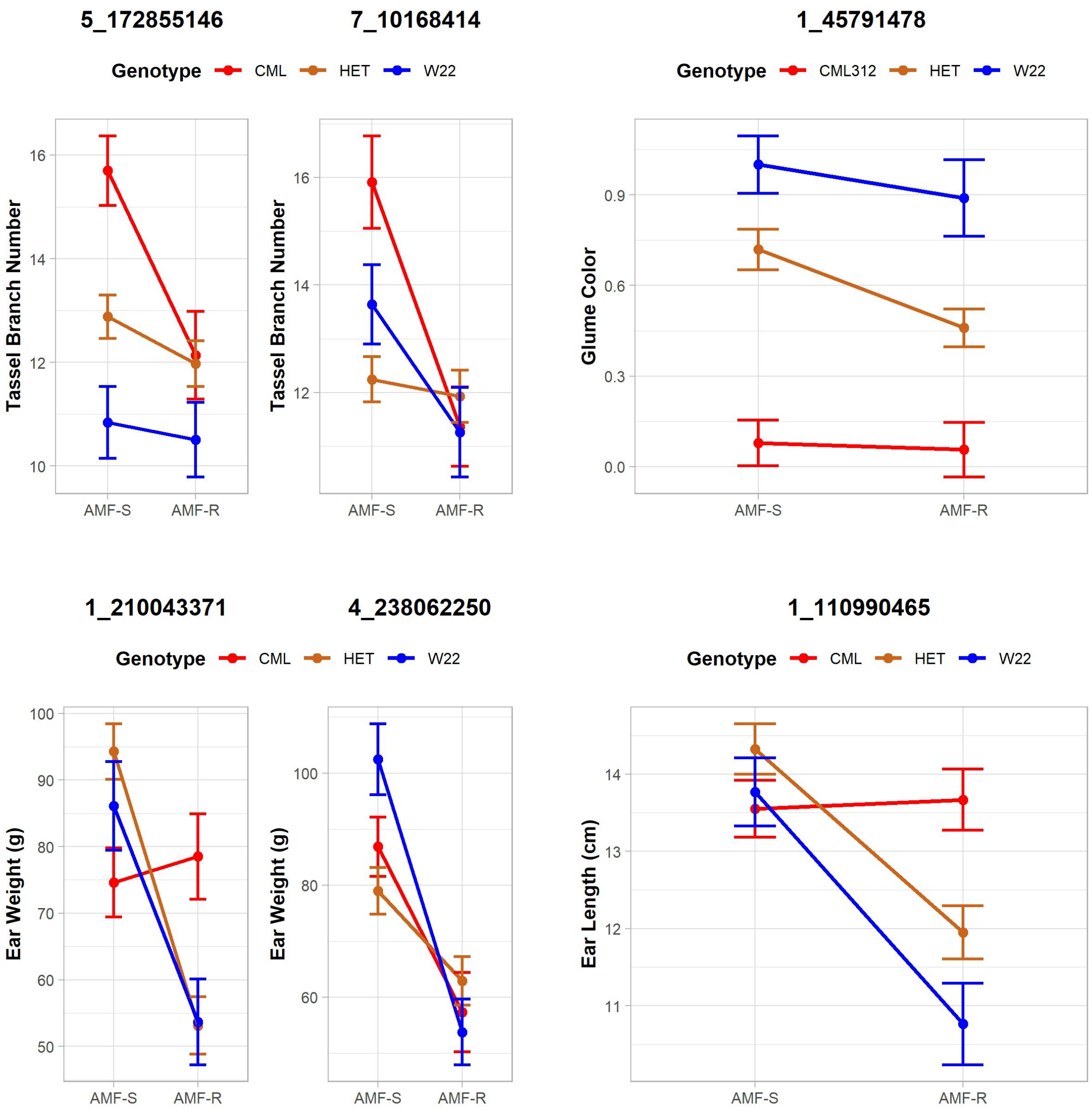

5_172855146
7_10168414
1_45791478
Genotype
CML
HET
W22
Genotype
CML312
HET
W22
Tassel Branch Number
16
14
12
10
Tassel Branch Number
16
14
12
Glume Color
0.9
0.6
0.3
0.0
AMF-S
AMF-R
1_210043371
4_238062250
1_110990465
Genotype
CML
HET
W22
Genotype
CML
HET
W22
Ear Weight (g)
100
90
80
70
60
50
Ear Weight (g)
100
80
60
Ear Length (cm)
14
13
12
11
AMF-S
AMF-R

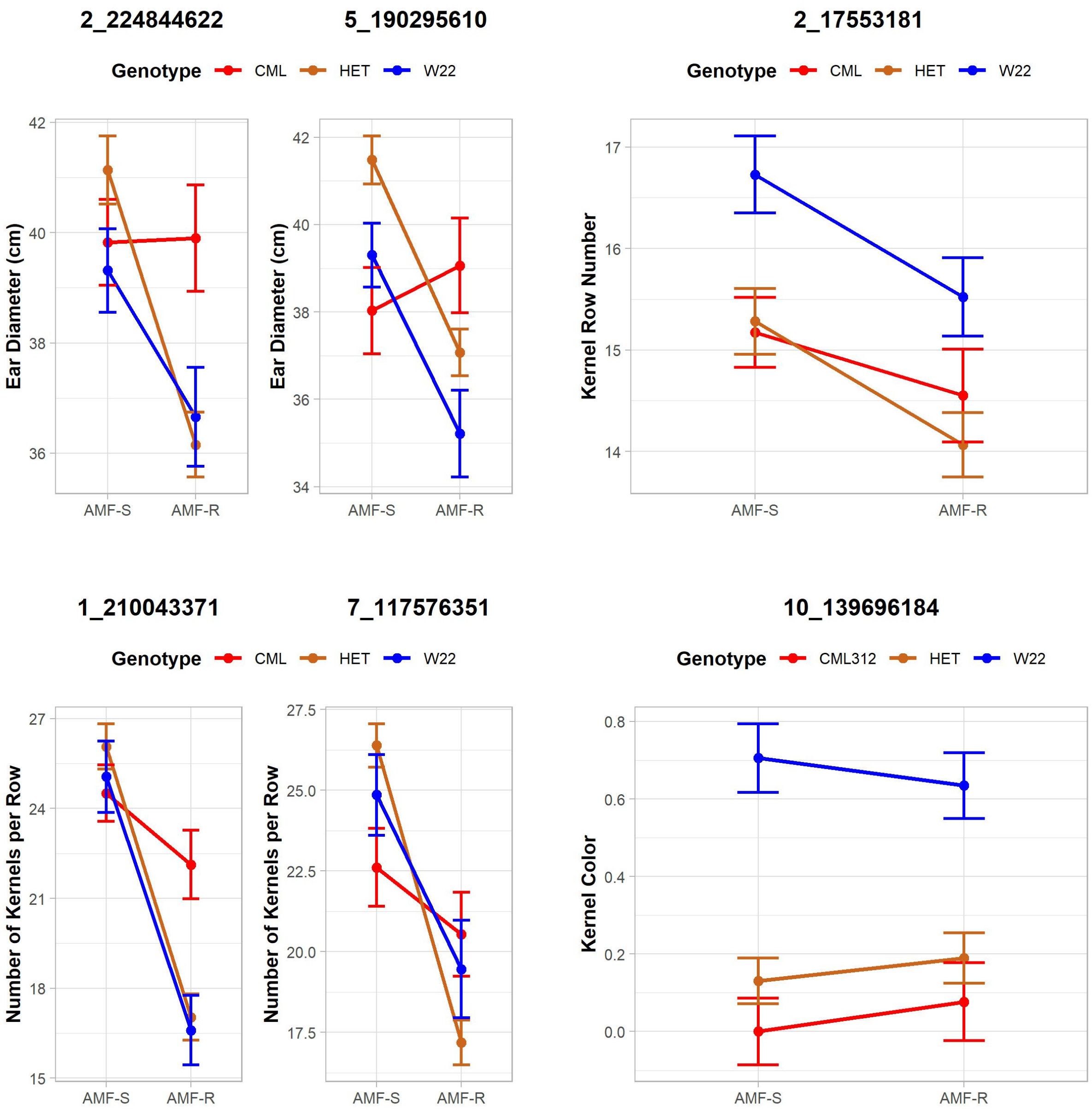
2_224844622
5_190295610
2_17553181
Genotype CML HET W22
Ear Diameter (cm)
Kernel Row Number
AMF-S
AMF-R
1_210043371
7_117576351
10_139696184
Genotype CML312 HET W22
Number of Kernels per Row
Kernel Color

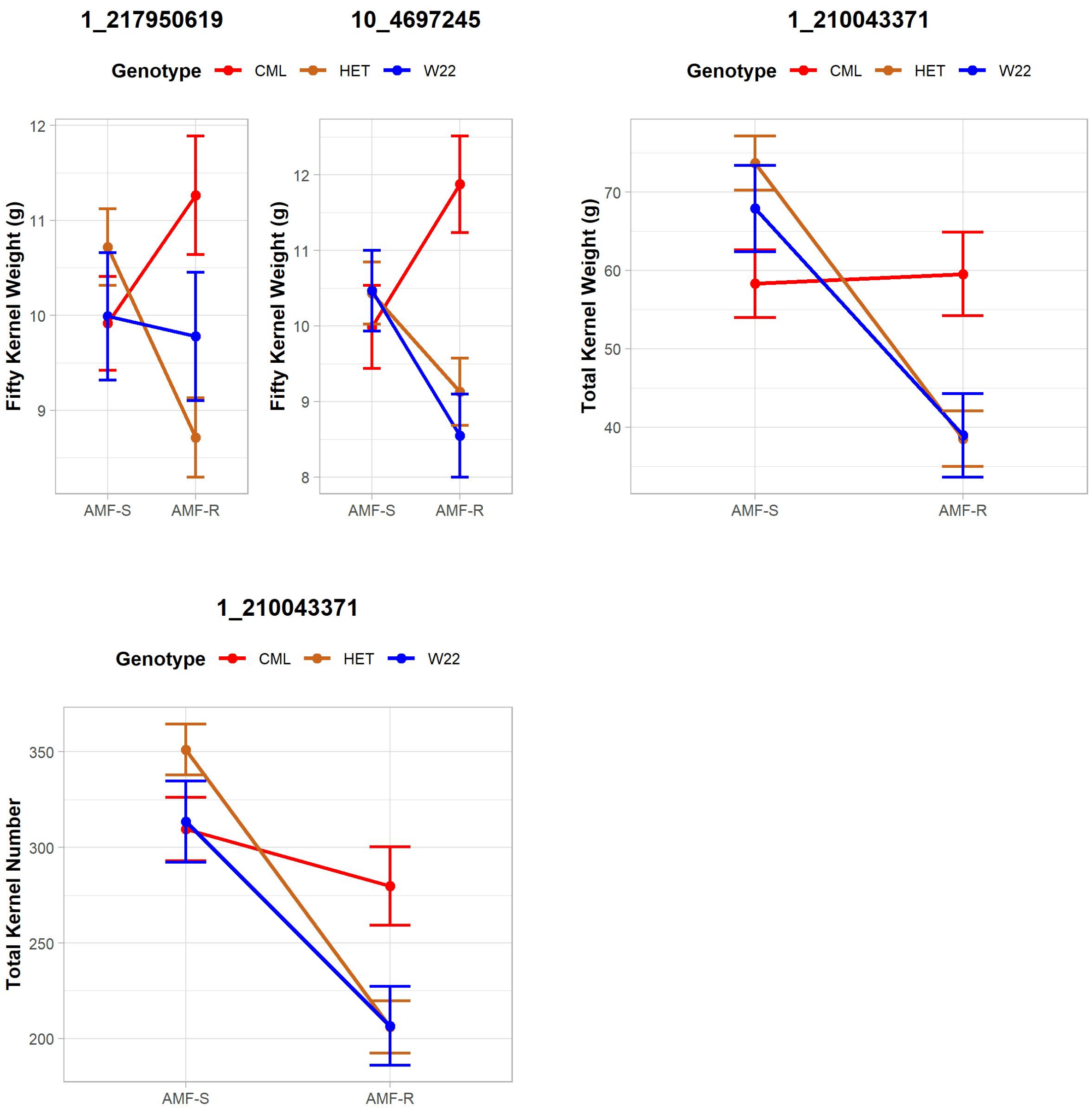


**Figures S14. Effect plots for major QTL detected in the analysis across susceptible (AMF-S) and resistant (AMF-R) families**. The title of the plot gives the marker nearest to the LOD peak, and the color of the line represents the genotype at the QTL.

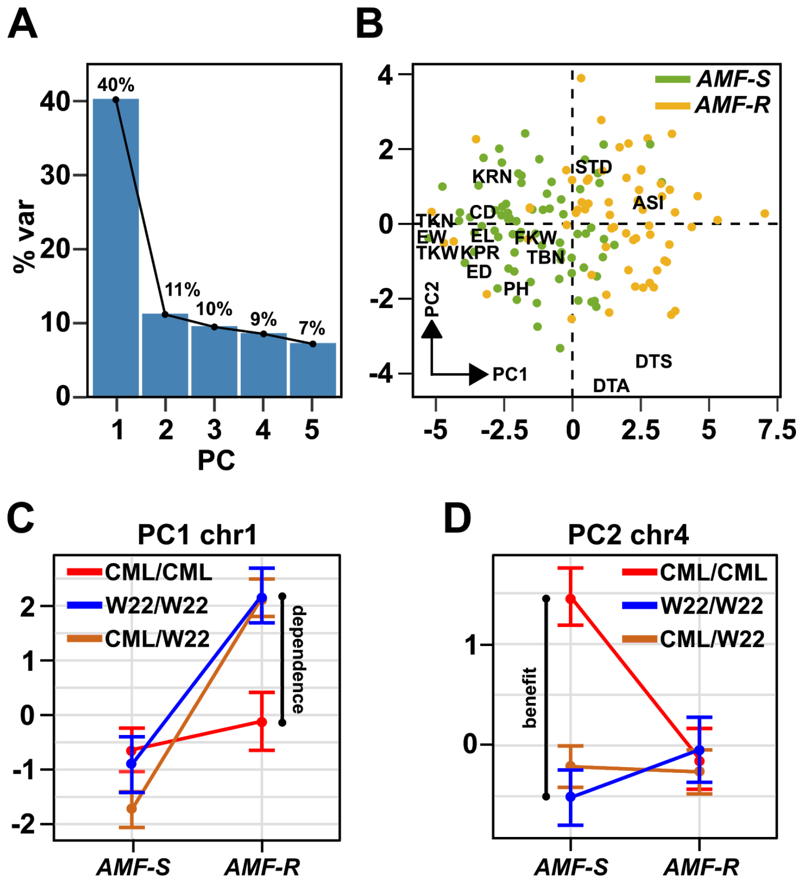


**Figure S15. Trait PCs capture variation in both dependence and benefit. A**, Phenotypic variation captured by the first five principal components (PCs). The first two principal components (PCs) explained more than 50% of the phenotypic variance. PC1 (40.1%) was primarily associated with ASI and maize yield-related traits, such as ear weight (EW), total kernel number (TKN), and total kernel weight (TKW). **B**, Loading of *AMF-S* and *AMF-R* families on PC1 and PC2. The position of trait names (as Table 1) with respect to the origin indicates contribution to each PC. **C, D,** effect plots for major QTL associated with PC1 and PC2, respectively. The PC1 QTL is conditional on *AMF-R*, indicating a difference in dependence. The PC2 QTL is conditional on *AMF-S*, indicating a difference in benefit.

### Supplementary Tables

**Table S1: Description of phenotypic traits measured in this study**

| Phenotypic traits | Description | Unit |
|---|---|---|
| DTA | Days to anthesis | Day |
| DTS | Days to silking | Day |
| ASI | Anthesis-silking interval | Day |
| PH | Plant height | cm |
| TBN | Tassel branch number | Count |
| EW | Ear weight | g |
| EL | Ear length | cm |
| ED | Ear diameter | cm |
| KRN | Number of kernel rows | Count |
| KPR | Number of kernels per row | Count |
| CD | Cob diameter | cm |
| 50GW | Weight of 50 kernels | g |
| TKW | Total kernel weight | g |
| TKN | Total kernel number | Count |

**Table S2. Broad sense heritability (H2) and percentage of phenotypic variance explained by *HUN* additive term (AMF), interaction between *HUN* and the QTLs (QTLxAMF) and QTL additive term (QTL) in the multi-qtl model for each trait .**

| Trait | H2 | AMF | QTLxAMF | QTL |
|---|---|---|---|---|
| STD | 41.6 | | | |
| DTA | 81.2 | 0.1 | 6.9 | 6.9 |
| DTS | 61.6 | .2 | 9.4 | 6 |
| ASI | 50.1 | 17.5 | 8.5 | 3.4 |
| PH | 68.2 | 39 | 6.6 | 4 |

| | | | | |
|---|---|---|---|---|
| TBN | 75.2 | 4.1 | 6.5 | 15.9 |
| EW | 54.6 | 31.2 | 7.2 | 1.7 |
| EL | 63.1 | 16.3 | 10 | 3.6 |
| ED | 56.8 | 25 | 6 | 5.4 |
| CD | 64.1 | | | |
| KRN | 62.8 | | | 11.1 |
| KPR | 55.4 | 43.7 | 6 | 3.5 |
| KC | 70.6 | | | 40.5 |
| GC | 79.4 | | | |
| FKW | 59.4 | 10.2 | 6.2 | 4.7 |
| TKW | 57.7 | 25.2 | 10.5 | 0.7 |
| TKN | 52.4 | 30.1 | 6.3 | 2 |
